# JUN mediates senescence and immune cell recruitment to prevent prostate cancer progression

**DOI:** 10.1101/2023.11.29.569178

**Authors:** Torben Redmer, Martin Raigel, Christina Sternberg, Roman Ziegler, Clara Probst, Desiree Lindner, Astrid Aufinger, Tanja Limberger, Karolina Trachtova, Petra Kodajova, Sandra Högler, Michaela Schlederer, Stefan Stoiber, Monika Oberhuber, Marco Bolis, Heidi A. Neubauer, Sara Miranda, Martina Tomberger, Nora S. Harbusch, Ines Garces de los Fayos Alonso, Felix Sternberg, Richard Moriggl, Jean-Philippe Theurillat, Boris Tichy, Vojtech Bystry, Jenny L. Persson, Stephan Mathas, Fritz Aberger, Birgit Strobl, Sarka Pospisilova, Olaf Merkel, Gerda Egger, Sabine Lagger, Lukas Kenner

## Abstract

**Background:** Prostate cancer develops through malignant transformation of the prostate epithelium in a stepwise, mutation-driven process. Although activator protein-1 transcription factors such as JUN have been implicated as potential oncogenic drivers, the molecular programs contributing to prostate cancer progression are not fully understood.

**Methods:** We analyzed JUN expression in clinical prostate cancer samples across different stages and investigated its functional role in a *Pten*-deficient mouse model. We performed histopathological examinations, transcriptomic analyses and explored the senescence-associated secretory phenotype in the tumor microenvironment.

**Results:** Elevated JUN levels characterized early-stage prostate cancer and predicted improved survival in human and murine samples. Immune-phenotyping of *Pten*-deficient prostates revealed high accumulation of tumor-infiltrating leukocytes, particularly innate immune cells, neutrophils and macrophages as well as high levels of STAT3 activation and IL-1β production. *Jun* depletion in a *Pten*-deficient background prevented immune cell attraction which was accompanied by significant reduction of active STAT3 and IL-1β and accelerated prostate tumor growth. Comparative transcriptome profiling of prostate epithelial cells revealed a senescence-associated gene signature, upregulation of pro-inflammatory processes involved in immune cell attraction and of chemokines such as IL-1β, CCL3 and CCL8 in *Pten*-deficient prostates. Strikingly, JUN depletion reversed both, senescence and senescence-associated immune cell infiltration and consequently accelerated tumor growth.

**Conclusions:** Our results suggest that JUN acts as tumor-suppressor and decelerates the progression of prostate cancer by transcriptional regulation of senescence- and inflammation-associated genes. This study opens avenues for novel treatment strategies that could impede disease progression and improve patient outcomes.

## Background

Prostate cancer (PCa) is one of the most frequently diagnosed malignancies in men worldwide [1]. Its significance lies not only in its prevalence but also in its potential to progress to aggressive forms that resist conventional treatments and lead to high mortality rates [2]. The complex molecular programs that determine the routes of PCa progression are still incompletely understood. On the molecular level, the dysregulation of the phosphoinositide 3-kinase (Pl3K) and androgen receptor (AR) pathways has been implicated in the pathology of PCa [3]. The constitutive activation of the Pl3K cascade, which is caused by mutations in the tumor-suppressor gene and Pl3K antagonist *Phosphate and tensin homologue* (*PTEN),* was identified in 20% of primary PCa tumors and represents a major oncogenic driver [4]. The current standard treatment for primary advanced-stage PCa is the administration of anti-androgens to deprive the tumor of dihydrotestosterone. PCa inevitably escapes androgen deprivation by relapsing into castration resistant PCa (CRPC), which is associated with loss of *PTEN* tumor-suppressor activity in 50% of cases. The characteristic dissemination of CRCP into local and distant regions such as bone, is correlated with poor survival [3–5].

In a previously described mouse model, the abrogation of *Pten* in prostate epithelium (PE) caused activation of a p53-mediated senescence program [6–8]. The emergence of senescence in cancer is considered a double-edged sword: it either confers anti-tumorigenic effects when originating from tumor cells or results in pro-tumorigenic outcomes when the tumor microenvironment (TME) is affected [9]. This phenomenon is mainly attributed to the induction of a senescence-associated secretory phenotype (SASP), characterized by the secretion of soluble signaling factors, proteases and extracellular matrix proteins [10]. In particular, pro-inflammatory cytokines such as IL-6, IL-1, CCL3 and CCL8 attract innate immune cells to the vicinity of the tumor site. As a collective, all components of SASP aid in creation of a pro-tumorigenic microenvironment and ultimately advance tumor progression depending on the tissue context. IL-6 and its downstream effector signal transducer and activator of transcription 3 (STAT3) are known to regulate apoptosis, angiogenesis, proliferation and differentiation, making them promising therapeutic targets in PCa [11]. However, our group has recently challenged active IL-6/STAT3 signaling as a tumor driver in PCa, as loss of *Stat3* unexpectedly resulted in increased tumor burden and was accompanied by a bypass of Pten-loss induced cellular senescence (PICS) in a *Pten*-deficient PCa mouse model [12,13].

Besides the hyperactivation of PI3K/AKT and amplification of AR signaling, other mechanisms driving the progression of PCa include the activation of activator protein-1 (AP-1) mediated gene expression [14]. AP-1 transcription factors (TF) such as JUN, were initially considered as proto-oncogenes [15] and deregulation of AP-1 family members was observed in several cancers [16]. Previous studies have suggested that JUN modulates hepatocellular tumorigenesis as a regulator of cell cycle genes and has co-activator and repressor functions in the regulation of AR in the prostate [17–19]. Recent evidence suggests tumor-suppressive functions for several members of the AP-1 TF family and their regulators [17,20]. For example, the JUN-activating JUN N-terminal kinase (JNK) has previously been identified as a potent tumor-suppressor in a murine PCa model [21]. JUNB, which is also activated by JNK has been associated with growth limiting properties in PCa and its activation may explain the mechanism of JNK’s tumor-suppression [22]. A recent study provides novel insights how the tumor-suppressive functions of AP-1 might be exerted, as JUN was particularly implicated as pioneering factor in bookmarking the enhancers of genes associated with the induction of the senescence program [23].

Here we investigated the role of *Jun* in a murine model of *Pten*-loss driven neoplasia of the PE and surveyed the consequence of JUN-deficiency in tumor development and senescence.

## Methods

### Mouse strains and animal work

To establish the PCa mouse model used in this study, we bred a *Pten* knockout prostate cancer mouse strain (*Pten^PE^*^Δ/Δ^) [24] with a *Jun*-floxed (*Jun^fl/fl^*) [25] mouse strain. The *Pten^PE^*^Δ/Δ^ mouse strain was originally established by crossing *Pten^Ex4/Ex5^*-floxed mice [26] and heterozygous transgenic *Probasin* (*Pb*) *Cre* mice [27]. *Pb Cre* transgenic mice express the Cre recombinase under the *Probasin* promoter restricted to PE cells of sexually mature mice [27]. To minimalize tumor burden for breeding animals, heterozygous *Pten^PEΔ/+^* males were used for breeding. The resulting genotypes of experimental animals are: *PbCre^+/+^*(*wildtype (wt)*), *PbCre^tg/+^*;*Jun*^fl/fl^ (*Jun^PE^*^Δ/Δ^); *PbCre^tg/+^*;*Pten*^fl/fl^ (*Pten^PE^*^Δ/Δ^); *PbCre^tg/+^*;*Jun*^fl/fl^;*Pten*^fl/fl^ (*Jun^PEΔ/Δ^;Pten^PEΔ/Δ^*). For all experiments, mice were sacrificed at 19-weeks of age, with the exception of animals used for the Kaplan-Meier survival analysis.

### Magnetic cell sorting, library preparation and RNA sequencing

The preparation of sequencing libraries and subsequent RNA sequencing (RNA-seq) was performed as previously described [28]. Briefly, prostates of 19-week-old mice were dissected, processed to yield a single cell suspension and EpCAM (CD326) positive cells were isolated by magnetic cell sorting (Magnisort®, Thermo Fisher Scientific) using anti-CD326-biotin (13– 5791-82, eBioscience). EpCAM positive cells were collected by centrifugation at 300 xg for 5 min at 4°C and stored at −80°C until further use. High-quality RNA, as assessed by 4200 TapeStation System (Agilent) was used for library preparation according to the manufacturer’s instructions.

### RNA sequencing data analysis

Single-end 75 bp reads sequencing of libraries was performed at CEITEC, Centre for Molecular Medicine (Brno, Czech Republic) as previously described [28]. Genes with a false discovery rate (FDR) FDR-adjusted p-value < 0.05 and log2 fold change ≥ 1 or ≤ −1 were considered significantly up- or downregulated.

### Histological staining

H&E, IHC and immunofluorescence (IF) stainings were performed on 2 µm sections of FFPE tissue. H&E staining was done according to routine diagnostic protocols. Details of IHC staining for the different markers are indicated in Supplementary Table 6 and all slides were counterstained with hematoxylin.

For the EpCAM IF staining, slides were dewaxed and heated in pH 6 citrate buffer. After blocking with 2% bovine serum albumin (Roth 8076.4), the slides were incubated in primary antibody (EpCAM, Elab Science, E-AB-70132, dilution 1:300) overnight. Next, slides were incubated for 1 hour at room temperature in secondary antibody (Goat anti-Rabbit IgG Alexa Fluor 488, Dilution 1:500) and stained with DAPI.

### Human tissue microarray analysis

The generation of human TMAs of healthy and tumor prostate tissues was previously described [33]. The TMAs were stained with an antibody for JUN (Supplementary Table 6) and analysed by trained pathologists. Staining was quantified by combining staining intensity with percentage of positive cells and divided in absent (0) or present (1) JUN expression. We then correlated the JUN status with BCR data and visualized the results in a Kaplan-Meier curve.

### Whole slide scan analysis

Analysis of IHC staining was performed with QuPath (version 0.3.2) [34]. First, regions of interest were annotated, excluding non-prostate tissue such as urethra, seminal vesicles and ductus deferens. Cell detection was performed with the StarDist extension [35] for the NIMP-R14 staining and the built-in watershed cell detection plugin for F4/80, CD79b, JUN and phosphorylated (p)STAT3. Parameters were chosen individually for each staining. Thereafter, smoothed features were calculated with a FWHM radius of 25 µm. The tissue was then classified into tumor/epithelium and stroma using an object classifier, trained individually for each staining. A threshold was set for the mean DAB optical density value, categorizing cells into positive or negative. For pSTAT3, multiple thresholds were set and cells were classified into 1*, 2* and 3* positive to calculate the H-score. The H-score was calculated by multiplying the percentage of cells by their respective intensity value and ranged from 1 to 300. Analysis was performed by a single investigator and evaluated by two independent pathologists. For quantification of Ki67 levels of tumor and non-tumor samples, we defined four circular regions of interest with a radius of 150 µm. Within each region, we manually counted the positive epithelial cells and used QuPath to detect the negative cells. Results shown are from the anterior prostate.

### Statistical analysis for immunohistochemistry

Measurements were exported as TSV files and imported into GraphPad PRISM (version 9.5.0). Significance was determined using an ordinary one-way ANOVA with Tukey’s multiple comparisons tests for 3 or more groups. Graphs were created and formatted in GraphPad PRISM.

### Protein extraction and immune blotting

Protein extraction from frozen prostate samples and immune blotting was performed as previously described [36]. Briefly, 15–20 µg of protein lysate were separated via SDS-PAGE, transferred onto nitrocellulose membranes (Amersham) and blocked with 5% milk in 1× TBS /0.1% Tween-20 or with 5% BSA in 1× TBS /0.1% Tween-20 for 1 h according to manufacturer’s antibody datasheets. Membranes were incubated with primary antibodies against pJUN^S73^ (CST 9164), JUN (CST 9165), pAKT^S473^ (CST 4060), AKT (CST 4691), EpCAM (Elab Science, E-AB-70132), β-ACTIN (CST 4967), NLRP3 (CST 15101), Pro-IL-1β (R&D Systems, AF-401-NA) and β-TUBULIN (CST 2146 and CST 2128) at 4°C overnight. TGX stain free technology (Bio-Rad), β-ACTIN or β-TUBULIN were used as loading controls.

## Results

### *JUN* levels discriminate progression states in prostate cancer

To clarify the role of AP-1 TFs in PCa progression, we investigated the level of the master factor JUN in tissue microarrays (TMA) of low and high progressive human prostate tumors by immunohistochemistry (IHC). We performed semi-quantitative analysis and categorized each tumor based on JUN levels ranging from 0 (absent) to 1 (present). Patients were divided into high (n=29 + 6 censored subjects) and low (n=32 + 8 censored subjects) JUN expression cohorts and correlated with biochemical recurrence (BCR) data. PCa progression is marked by histological changes of the tumor architecture and is categorized by Gleason scoring [38]. We observed high JUN protein abundance in primary tumors (low Gleason) while JUN levels were reduced in advanced tumor stages (high Gleason) (Fig. 1a). The correlation between JUN protein and patient’s BCR status revealed a significantly (p=1.8e-02) diminished BCR free survival in patients with low JUN (Supplementary Fig. 1a), whereas high JUN levels were associated with increased survival probability. We next mined a publicly available transcriptome dataset ([39]; n=140) and stratified PCa patients into high-risk and low-risk groups as defined by the prognostic index and characterized by a significant difference in survival (p=4e-04) (Supplementary Fig. 1b). We correlated the high- and low-risk groups with *JUN* mRNA and found significantly (p=1.3e-30) higher *JUN* among low-risk patients compared to the high-risk group (Supplementary Fig. 1c). In a second dataset ([40]; n=333), we applied the KMplot tool to assess the relapse-free survival (RFS) and detected significantly (p=8.4e-03) decreased survival probability in patients showing low *JUN* expression (*JUN^low^*) compared to increased survival of patients expressing high *JUN* (*JUN^high^*) (Fig. 1b). In summary, the results from public databases confirmed the initial findings of the TMA analysis.

**Figure 1:**
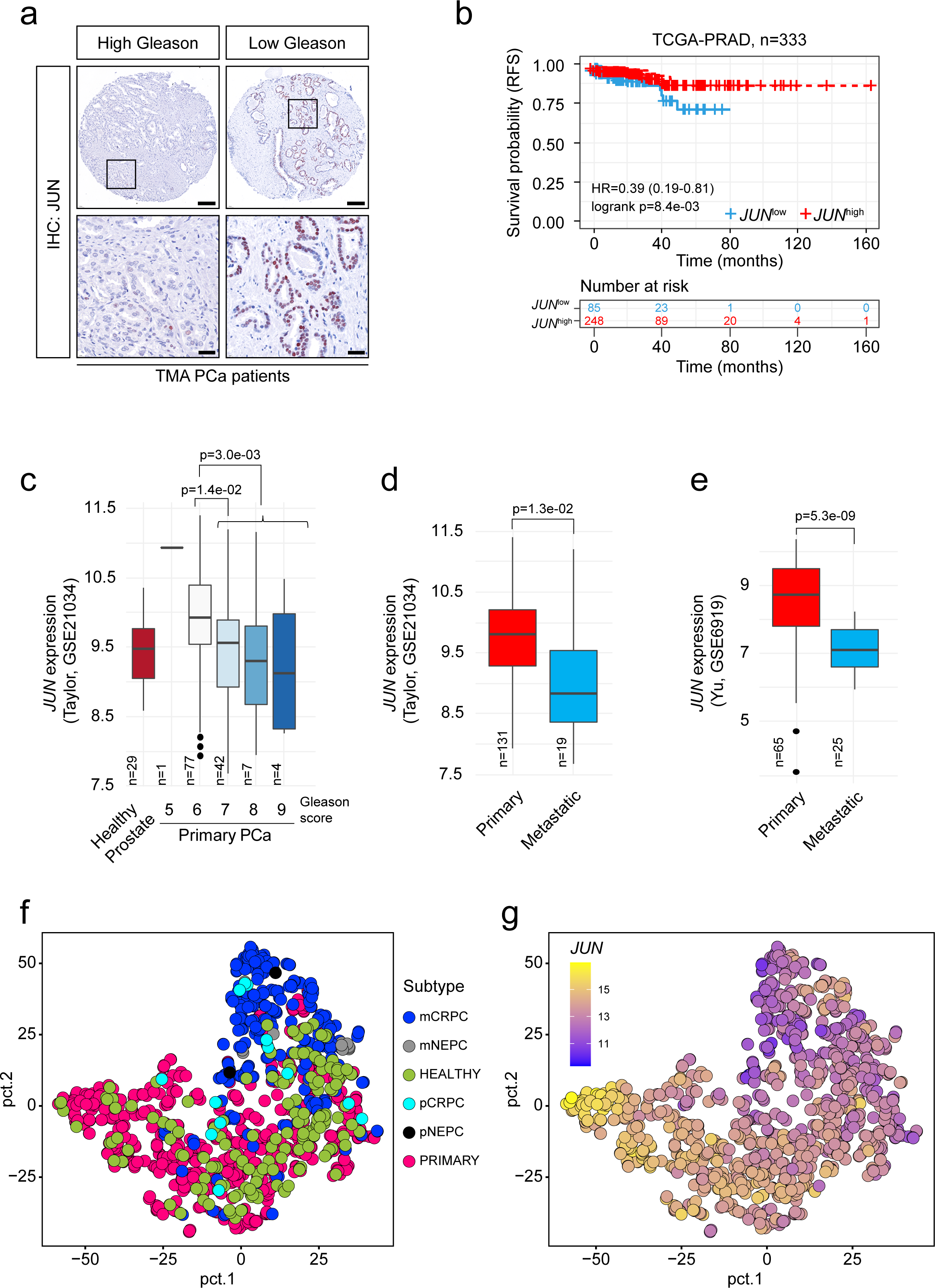
JUN levels are correlated with prostate cancer progression stages. a) Representative immunohistochemistry (IHC) images of tissue microarrays (TMAs) investigating human prostate tumors (n=60) with high or low Gleason scores stained for JUN protein. Scale bars indicate 150 µm (top row) and 30 µm (bottom row), images are presented in 16.8x (top row) and 80.0x magnification (bottom row). The area used for the higher magnification is indicated by the rectangle. b) Kaplan-Meier survival analysis of TCGA-PRAD [40] tumors (n=333) indicating a low hazard ratio (HR<1) and reduced risk of relapse-free survival (RFS) of *JUN^high^* (red line) compared to *JUN^low^* (blue line) tumors. Statistical testing was done with a logrank test. c) *JUN* mRNA levels in high (Gleason score ≥ 7) and low (Gleason score < 7) grade human prostate tumors. Data were retrieved from [39]. Significance was determined by an unpaired, two-sided t-test or one-sided Anova. d) High and low *JUN* levels significantly (p=1.3e-02) discriminate primary (n=131) (red) and metastatic (n=19) (blue) prostate tumors. Data were retrieved from [39]. e) High and low *JUN* levels significantly (p=5.3e-09) discriminate primary (n=65) (red) and metastatic (n=25) (blue) prostate tumors. Data were retrieved from [41]. f) Principal component analysis (PCA) of prostate tumors of different developmental stages comprising normal prostate tissue, primary tumors and primary (p) and metastatic (m) CRPC and NEPC tumors. Datasets from [42]. g) Overlay of *JUN* expression with PCA clustering from f). *JUN* levels are color coded from high expression (yellow) to low expression (blue). Significances in c-e were determined by an unpaired, two-sided t-test.

To explore *JUN* levels in advanced stages of PCa, we used the Taylor dataset [39], comprising primary tumors of different progression stages and Gleason scores (n=131) as well as healthy prostate tissue (n=29). Compared to healthy tissue, we observed higher levels of *JUN* in early disease stages with Gleason scores 5-6 and significantly decreased expression of *JUN* in high grade tumors (p=3e-03; Gleason scores 7-9) (Fig. 1c). Concordantly, *JUN* was highly expressed in primary tumors (n=131; n=65) but significantly lower expressed in PCa metastases (n=19; n=25) as observed in two independent datasets (Figs. 1d-e; p=1.3e-02; [39]; p=5.3e-09; [41]). We next investigated levels of JUN dimerization partners, JUNB and FOS and observed a comparable regulation of both (Supplementary Figs. 1d-e). Metastatic CRPC and neuroendocrine PCa (NEPC) present aggressive tumor subtypes that emerge under androgen deprivation therapy and are associated with poor prognosis. We compared levels of *JUN* and its related TFs *FOS* and *JUNB* in primary (n=715) and metastatic (n=320) PCa [42], including CRPC and NEPC (Figs. 1f-g, Supplementary Fig. 1f). The tumor-subtype and stage-dependent expression of *JUN* was highly significant when comparing healthy and primary (p=2.8e-05), primary and metastatic CRPC (p=2.6e-43) and primary and metastatic NEPC (p=5.3e-04) (Supplementary Fig. 1g), suggesting JUN as a potential marker of aggressive subtypes of PCa. In addition, our survey revealed higher levels of *JUN* in primary PCa than healthy prostates (Supplementary Figs. 1g-h), suggesting a gradual change of *JUN* levels in PCa development and progression. Our data implicate that JUN and other AP-1 factors except ATF2, may act as suppressors rather than drivers of PCa which was reflected by hazard ratios (HR) calculated from BCR (Supplementary Fig. 1i).

### Genetic depletion identifies a tumor-suppressive role of JUN in prostate cancer development

To elucidate the mechanistic role of JUN in PCa development, we employed a well-established murine model of PCa [43] (Fig. 2a), harboring floxed exons 4 and 5 of *Pten* [26]. The homozygous deletion of murine *Pten* via the *Probasin (Pb)* Cre recombinase [27] mirrored 20% of all primary human PCa cases with homozygous loss of *Pten* (Fig. 2a, white mouse). The PE of homozygous mutants developed hyperplasia that progressed into prostate adenocarcinoma between 12 and 29 weeks of age [43]. We inter-crossed a floxed *Jun* mouse strain where the sole exon is flanked by loxP sites [25] (Fig. 2a, grey mouse) to generate 4 individual genotypes. This enabled comparison of prostate tissue of *wildtype (wt)* mice to either *Jun* (*Jun*^PEΔ/Δ^), *Pten* (*Pten*^PEΔ/Δ^) or *Jun*/*Pten* (*Jun*^PEΔ/Δ^; *Pten*^PEΔ/Δ^) double knockout mice (Fig. 2a, colored F1 mice). We examined protein extracts of whole prostates and observed a significant increase in levels of phosphorylated (S73) and total JUN in *Pten^PEΔ/Δ^*, whereas notable JUN expression was absent in *wt* prostates (Fig. 2b). We also confirmed efficient Cre-mediated deletion of *Jun* alone (*Jun*^PEΔ/Δ^) and in combination with *Pten* (*Jun*^PEΔ/Δ^; *Pten*^PEΔ/Δ^) (Fig. 2b). As a verification of functional *Pten* deletion, we detected robust activation of the PI3K/AKT pathway in *Pten*^PEΔ/Δ^ and *Jun*^PEΔ/Δ^; *Pten*^PEΔ/Δ^ mice as assessed by analysis of phosphorylated AKT levels (Fig. 2b).

**Figure 2:**
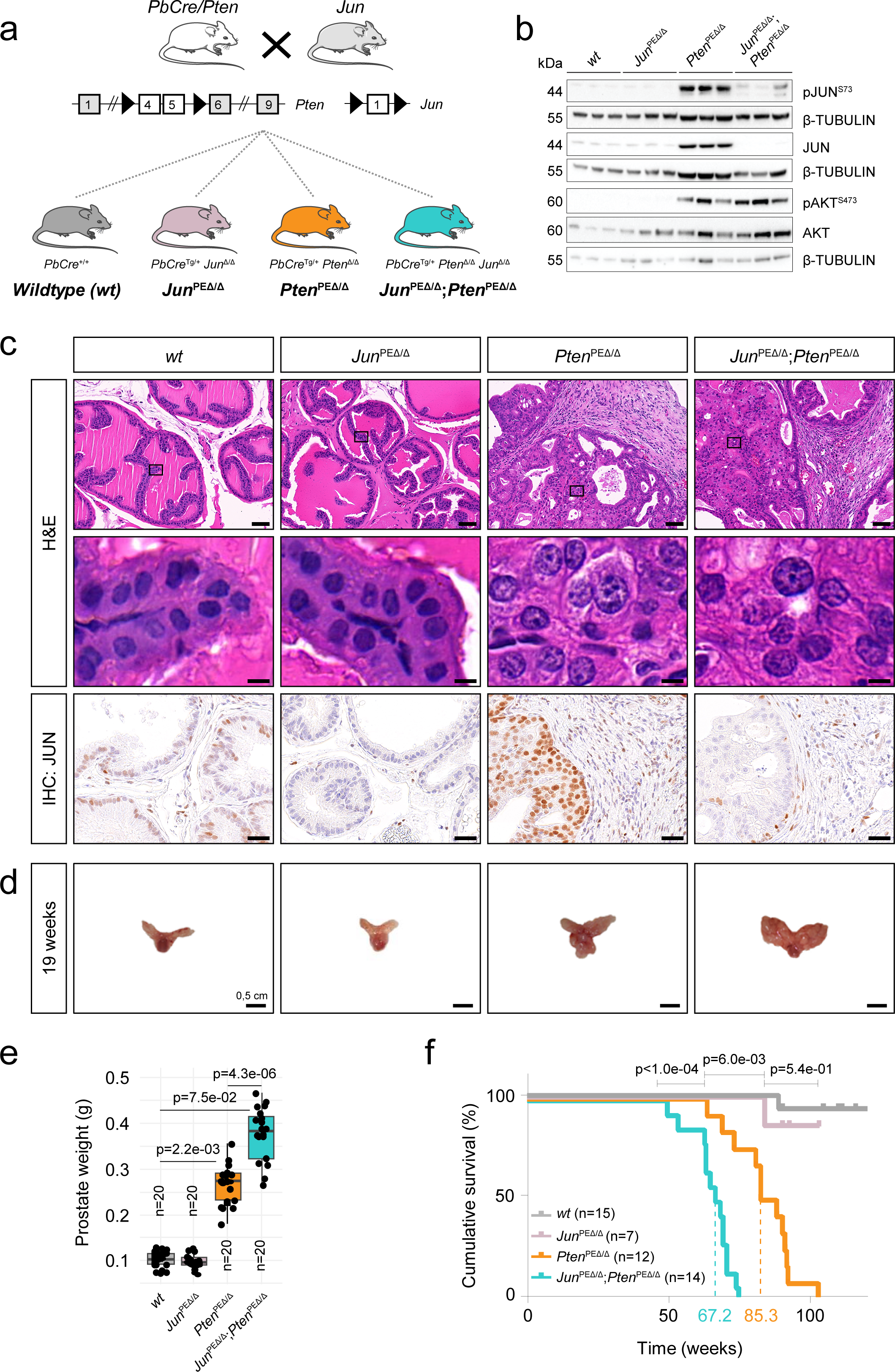
*Jun*-deficiency fosters the progression of *Pten*-loss induced tumors. a) Top: Schematic representation of mouse models used in the study. Homozygous loss of *Pten* or *Jun* was achieved by a *Probasin* promoter-controlled Cre recombinase (*PbCre*)-mediated ablation of floxed exons 4 and 5 (*Pten*) or exon 1 (*Jun*). Bottom: established and investigated genetic models. Wildtype (*PbCre^+/+^*; *wt*) and mice with single knockout of *Pten* (*PbCre ^tg/+^*; *Pten^PE^*^Δ/Δ^) and *Jun* (*PbCre ^tg/+^*; *Jun^PE^*^Δ/Δ^) were compared with double knockout (*PbCre ^tg/+^*; *Jun^PEΔ/Δ^;Pten^PEΔ/Δ^*). PE= prostate epithelium; tg = transgene; Δ = knockout. b) Western blot analysis of phosphorylated (pJUN^S73^ and pAKT^S473^) and total JUN and AKT. β-TUBULIN served as loading control. Protein lysates of entire organs (n=3 biological replicates) from 19-week-old *wt*, *Pten^PE^*^Δ/Δ^, *Jun^PEΔ/Δ^* and *Jun^PEΔ/Δ^;Pten^PEΔ/Δ^*were investigated. c) Top row: H&E stainings of 19-week-old *wt*, *Pten^PE^*^Δ/Δ^, *Jun^PEΔ/Δ^* and *Jun^PEΔ/Δ^;Pten^PEΔ/Δ^* prostates. Scale bars indicate 60 µm (top row) and 2 µm (second row), images are presented in 40.0x (top row) and 600.0x magnification (second row). Black rectangles represent the area used for the zoom image below. Bottom row: IHC with an antibody against JUN in 19-week-old prostates of all four experimental groups. Scale bars indicate 30 µm; images are presented in 100.0x magnification. d) Macroscopic images of 19-week-old dissected prostates of *wt*, *Pten^PE^*^Δ/Δ^, *Jun^PEΔ/Δ^* and *Jun^PEΔ/Δ^;Pten^PEΔ/Δ^*. e) Box plot showing the weights of prostates in grams between *wt*, *Pten^PE^*^Δ/Δ^, *Jun^PEΔ/Δ^* and *Jun^PEΔ/Δ^;Pten^PEΔ/Δ^* 19-week-old animals (n=20). Significance was determined with an unpaired, two-sided t-test. f) Kaplan-Meier survival analysis of *wt*, *Pten^PE^*^Δ/Δ^, *Jun^PEΔ/Δ^*and *Jun^PEΔ/Δ^;Pten^PEΔ/Δ^* animals. Biological replicates are indicated and the cumulative survival (%) is shown. Statistical significance was calculated with a logrank test.

To investigate the morphological architecture of prostates upon *Jun* deletion in the PCa mouse model, we analyzed histological sections by hematoxilin and eosin (H&E) staining (Fig. 2c, top panel). Both *wt* and *Jun^PEΔ/Δ^*animals showed physiological growth patterns and morphology, characteristic for the respective prostate lobes. In *Pten^PEΔ/Δ^* and *Jun*^PEΔ/Δ^; *Pten*^PEΔ/Δ^ prostates, we observed hyperplastic epithelium growing in cribriform patterns into the lumen. Both groups showed anisocytosis, anisokaryosis and alterations in nucleus-to-cytoplasmic ratios, but largely without invasion of the stroma. We were unable to detect significant differences between *Pten^PEΔ/Δ^* and *Jun*^PEΔ/Δ^; *Pten*^PEΔ/Δ^ prostates regarding the grading of the proliferative lesions (data not shown).

Next, we analyzed JUN levels in prostates of all genotypes. Supporting our immunoblot results, IHC revealed increased levels of total JUN predominantly in the PE of *Pten^PEΔ/Δ^* mice and absence in epithelial cells of *Jun*^PEΔ/Δ^ and *Jun*^PEΔ/Δ^; *Pten*^PEΔ/Δ^ (Fig. 2c, bottom panel). We assessed the effects of *Jun* deficiency on tumor burden and survival by morphological and survival analyses. Macroscopically, prostates from *Pten^PEΔ/Δ^* and *Jun*^PEΔ/Δ^; *Pten*^PEΔ/Δ^ mice were notably enlarged as compared to *wt* or *Jun*^PEΔ/Δ^ prostates (Fig. 2d). This finding was corroborated by prostate weight analysis (Fig. 2e). The additional deletion of *Jun* on the *Pten*-deficient background resulted in even higher prostate weights, hinting at JUN’s potential function as a tumor-suppressor in murine PCa development. We performed a Kaplan-Meier survival analysis where overall survival or the occurrence of the discontinuation criteria according to the guidelines of the 3Rs principles were defined as the endpoint of the experiments (Fig. 2f) [44]. We observed comparable survival probabilities of *wt* and *Jun^PEΔ/Δ^* mice (p=5.4e-01) but a significantly decreased survival of *Pten^PEΔ/Δ^* (mean survival 85.3 weeks, p=6e-03) as compared to *wt* mice. Remarkably, the survival of *Pten^PEΔ/Δ^* mice was significantly (p<1e-04) reduced by the additional deletion of *Jun*. *Jun*^PEΔ/Δ^; *Pten*^PEΔ/Δ^ mice exhibited a mean survival of 67.2 weeks. We therefore conclude that *Jun*-deficiency alone is not sufficient to induce prostate tumorigenesis, but causes a significant increase in tumor burden and a significant reduction in overall survival in combination with *Pten* knockout. The results of our murine PCa model reinforce our observations from human PCa samples, suggesting that JUN acts as a tumor-suppressor in PCa.

To determine whether aberrant cellular proliferation contributes to enhanced tumor growth in *Jun*^PEΔ/Δ^; *Pten*^PEΔ/Δ^-deficient prostates, we assessed the number of Ki67^+^ epithelial cells by IHC. Although we noticed higher Ki67 levels in *Jun*^PEΔ/Δ^; *Pten*^PEΔ/Δ^ tumors by trend, the difference was not significant (p=1.3e-01) when compared to *Pten^PEΔ/Δ^*prostates (Supplementary Fig. 2a). This indicates that proliferation may not be the primary biological process influenced by JUN during PCa progression.

### Transcriptome profiling reveals JUN-mediated alterations in senescence and immune response

To elucidate the tumor cell-specific molecular programs regulated by JUN *in vivo*, we performed transcriptome profiling of PE cells across all four experimental murine groups (Fig. 2a). To obtain a homogenous epithelial fraction, we enriched prostate lysates for the Epithelial cell adhesion molecule (EpCAM) showing a uniform expression in PE cells (Fig. 3a, Supplementary Fig. 3a) via magnetic cell separation [28] (Fig. 3b, Supplementary Fig. 3b). The correlation analysis revealed high congruence between *Jun*^PEΔ/Δ^; *Pten*^PEΔ/Δ^ and *Pten^PEΔ/Δ^* tumor and *wt* and *Jun^PEΔ/Δ^* samples (Fig. 3c).

**Figure 3:**
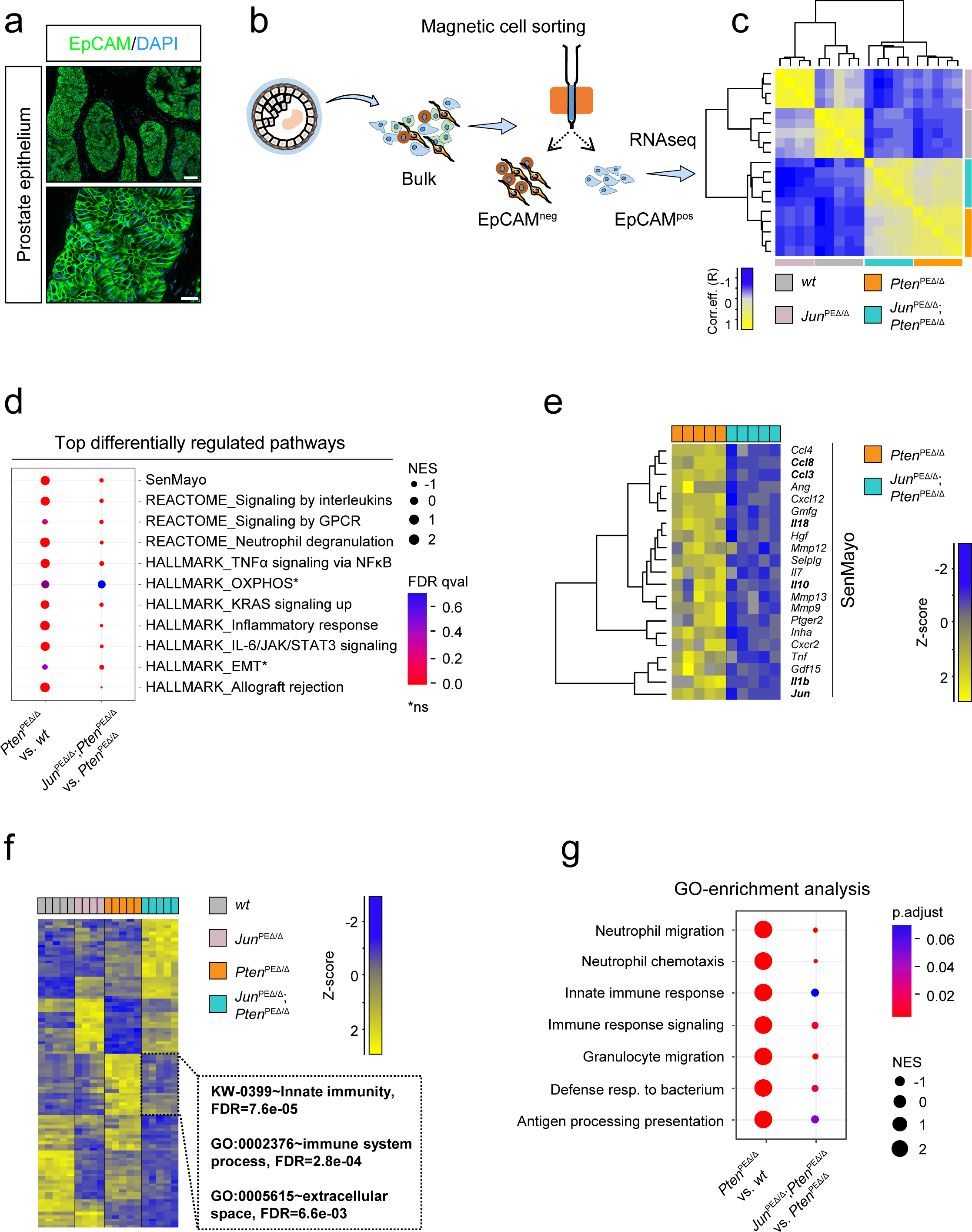
Transcriptome profiling of genetic models reveals a JUN-dependent regulation of innate immunity. a) Representative immunofluorescence (IF) image of a *wt* murine prostate for the epithelial marker EpCAM (green). DAPI (blue) is shown as a nuclear stain. Top image: 40.0x magnification, scale bar represents 60 µm; Bottom image: 147.5x magnification, scale bar represents 20 µm. b) Overview of sample preparation for transcriptome profiling of *wt*, *Pten^PE^*^Δ/Δ^, *Jun^PEΔ/Δ^* and *Jun^PEΔ/Δ^;Pten^PEΔ/Δ^* prostate samples of 19-week-old animals. An antibody against the epithelial marker EpCAM was used to separate single cell suspensions of minced and digested prostates into EpCAM positive (pos) and negative (neg) fractions by magnetic cell sorting. EpCAM^pos^ cells were used for RNA-seq expression profiling. c) Heat map showing correlation analysis of tumor samples described in b) regarding global similarity of samples. The pearson correlation coefficient (R) is shown (color coded). d) Gene onthology (GO)-enrichment analysis of differentially expressed genes (DEGs) showing the top differentially regulated pathways between *Pten^PE^*^Δ/Δ^ and *Jun^PEΔ/Δ^;Pten^PEΔ/Δ^*. Significance as shown by FDR is color coded, enriched (positive normalized enrichment score (NES)) or depleted (negative NES) processes are indicated. Asterisk represents non-significant pathways (ns). e) Heat map showing SenMayo genes most significantly (p≤1e-02) regulated among *Pten^PEΔ/Δ^* and *Jun^PEΔ/Δ^;Pten^PEΔ/Δ^*prostates. f) Heat map representation of *wt*, *Pten^PE^*^Δ/Δ^, *Jun^PEΔ/Δ^* and *Jun^PEΔ/Δ^*;*Pten^PEΔ/Δ^*samples showing DEGs. “Innate immunity”, FDR=7.64e-05; “Immune system”, FDR=2.77e-04 and “Extracellular space”, FDR=6.60e-03 related processes most discriminated the groups. Genotypes and expression levels are color coded. g) GO-enrichment analysis of DEGs showing the regulation of innate immune cells such as neutrophil granulocytes. Significance as shown by p-value is color coded, enriched (positive NES) or depleted (negative NES) processes are indicated. Shown are the signaling pathways enriched in *Pten^PE^*^Δ/Δ^ tumors compared to *wt* (left side) and *Jun^PEΔ/Δ^*;*Pten^PEΔ/Δ^*tumors compared to *Pten^PE^*^Δ/Δ^ (right side).

We next performed a comparative analysis of *Jun*^PEΔ/Δ^; *Pten*^PEΔ/Δ^ and *Pten^PEΔ/Δ^* prostate samples to discern JUN-dependent programs potentially contributing to PCa formation. Our survey revealed 1706 (p.adjust<5e-02) DEGs with top 102 genes being up- (log_2_fold change≥ 1) and top 91 genes downregulated (log_2_fold change≤ −1; Supplementary Table 1). DAVID analysis of top genes revealed increased “innate immunity” and “immune system processes” but decreased secretory-, extracellular matrix- and immune-related processes. Notably, *Jun* ranked among the top 10 downregulated genes confirming the successful knockout in epithelial cells (Supplementary Table 1). Gene set enrichment analysis (GSEA) revealed immune system-related processes, IL-6/STAT3 signaling and senescence-associated gene signatures among the most enriched processes in *Pten^PEΔ/Δ^* prostates which were significantly depleted in *Jun^PEΔ/Δ^*; *Pten^PEΔ/Δ^* (Fig. 3d). Our previous work suggested that activation of IL-6/STAT3 signaling and of the downstream acting p19^ARF^–MDM2–p53 axis contributed to senescence in *Pten^PEΔ/Δ^* prostates [12]. We therefore investigated the enrichment level of different senescence signatures including “oncogene-induced senescence” (OIS), “SASP” signatures and the novel “SenMayo” gene signature, consisting of 125 previously identified senescence/SASP-associated factors. SenMayo genes are transcriptionally regulated by senescence and allow identification of senescent cells across tissues [30]. We found significant (qval= 5.09e-04) enrichment of SASP in *Pten^PEΔ/Δ^* as compared with *wt* prostates (Supplementary Table 2). SenMayo genes were significantly (qval=2.40e-02) enriched in *Pten^PEΔ/Δ^* prostates and depleted (qval=2.64e-02) in *Jun^PEΔ/Δ^*; *Pten^PEΔ/Δ^*tumors (Fig. 3d). Among the depleted SenMayo genes in *Jun*-deficient *Pten^PEΔ/Δ^*prostates, we identified chemokines such as *Ccl3, Ccl4 and Ccl8*, along with pro-inflammatory cytokines such as *Il1b* and *Tnfa* (Fig. 3e). To further investigate the JUN-dependent regulation of senescence in *Pten*-deficient murine prostates, we stained formalin-fixed paraffin embedded (FFPE) material with the senescence marker p16^INK4A^ (Supplementary Fig. 3c). Although we did not observe differences in the amount of p16^INK4A^ positive cells between *Pten^PEΔ/Δ^* and *Jun^PEΔ/Δ^*; *Pten^PEΔ/Δ^*tumors, we found significant changes in staining patterns. While we detected prominent nuclear staining in *Pten^PEΔ/Δ^* samples, *Jun^PEΔ/Δ^*; *Pten^PEΔ/Δ^* revealed predominantly cytoplasmic localization, hinting at a potential inactivation of p16^INK4A^ via nuclear export [45]. These results implicate the establishment of a senescent phenotype in *Pten*-deficient prostates which is impeded upon additional *Jun* deletion.

As our results suggest JUN-dependent activation of the IL-6/STAT3 axis and our previous study connected loss of activated STAT3 in *Pten*-deficient PCa to increased tumor burden via disruption of senescence [12], we sought to analyze STAT3 tyrosine 705 (Y705) phosphorylation (pSTAT3^Y705^) in the *Jun*-deficient background. We indeed detected reduced levels of pSTAT3^Y705^ in both stroma (p=5.0e-04) and epithelial cells (p<1.0e-04) of *Jun*^PEΔ/Δ^; *Pten*^PEΔ/Δ^ compared to *Pten^PEΔ/Δ^* tumors (Supplementary Fig. 3d, Supplementary Fig. 3e, upper panel) while total STAT3 levels remained constant (Supplementary Fig. 3e, lower panel). Our findings provide evidence that loss of JUN accompanied by reduced activation of STAT3 bypasses the senescence mechanism and subsequently amplifies the tumor load in *Jun*^PEΔ/Δ^; *Pten*^PEΔ/Δ^ animals. We suggest JUN as a regulator of STAT3-mediated senescence in PCa *in vivo*, reinforcing JUN’s proposed function as a pioneering factor of senescence [23].

### Jun deficiency in the PCa mouse model leads to downregulated chemotaxis of innate immune cells

We next compared *Jun*^PEΔ/Δ^; *Pten*^PEΔ/Δ^ and *Pten^PEΔ/Δ^* prostate samples to uncover additional JUN-dependent biological processes involved in PCa formation. A stringent selection identified ∼100 significantly deregulated genes (padj≤1.0e-03, FClog2 ≤ −1.2; n=59/ FClog2 ≥ 1.2; n=46; Supplementary Table 3) and uncovered innate immunity and other immune system-related processes as most distinguishing between *Jun*^PEΔ/Δ^; *Pten*^PEΔ/Δ^ and *Pten^PEΔ/Δ^* prostate tumors (Fig. 3f). Amongst the innate immunity and immune system cluster, gene onthology (GO)-enrichment analysis indeed confirmed immune system-related signatures that were activated in *Pten^PEΔ/Δ^* and significantly reduced by *Jun*-deficiency (Fig. 3g). Innate immunity-related processes are complex and encompass more than 2,000 publicly available human and mouse annotated genes [31]. We defined a core immunity-related signature by GSEA applying 645 innate immunity-related genes and investigated the enrichment specifically in *Pten^PE^*^Δ/Δ^ prostates. The analysis revealed 111 genes, of which 26 were significantly (p<1.0e-03) differentially expressed between *Pten^PE^*^Δ/Δ^ and *Jun*^PEΔ/Δ^; *Pten*^PEΔ/Δ^ prostates (Fig. 4a, top panels, Supplementary Fig. 4a, Supplementary Table 4). Using the “Hallmark Inflammatory response” signature, we uncovered a similar pattern as the majority of genes from both signatures were significantly (p<1.0e-03) elevated in *Pten^PE^*^Δ/Δ^ and depleted in *Jun*^PEΔ/Δ^; *Pten*^PEΔ/Δ^ prostates (Fig. 4a, bottom panels, Supplementary Fig. 4a). Hence, the homozygous loss of *Pten* was accompanied by inflammation and inflammatory response likely driven by increased levels of *Il1b*, *Nlrp3* and chemokines such as *Ccl5*.

**Figure 4:**
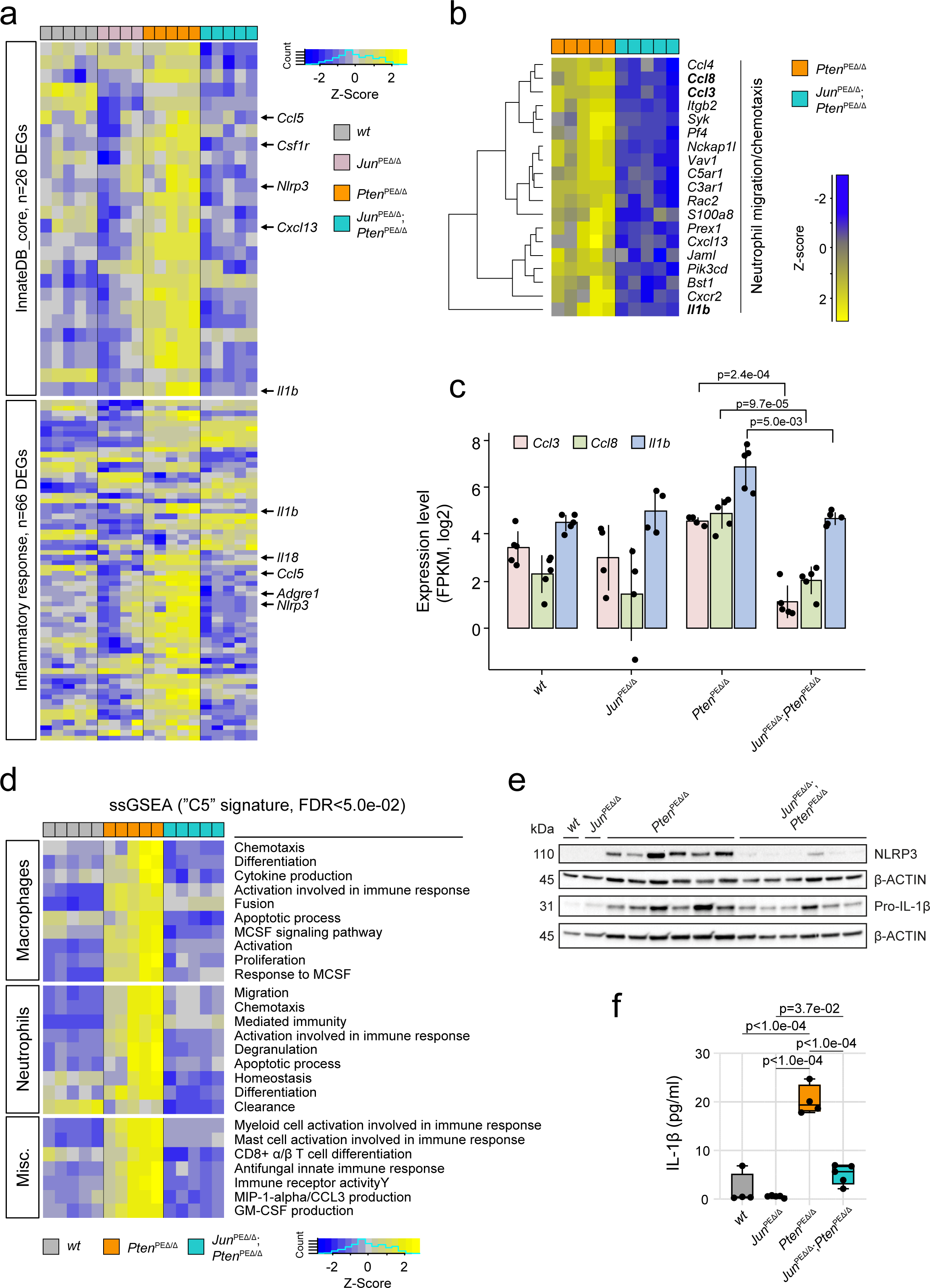
JUN expression determines the level of immune cell infiltration of *Pten*-loss driven tumors. a) Heat map showing JUN-dependent regulation of genes related to innate immunity (upper panel) and inflammatory response (lower panel) in *wt*, *Jun^PE^*^Δ/Δ^, *Pten^PE^*^Δ/Δ^ and *Jun^PEΔ/Δ^*;*Pten^PEΔ/Δ^*prostates. JUN-dependent core factors such as *Il1b*, *Nlrp3* and *Ccl5* are highlighted. b) Heat map presenting the JUN-dependent regulation of genes involved in migration and chemotaxis of neutrophil granulocytes in *Pten^PE^*^Δ/Δ^ and *Jun^PEΔ/Δ^*;*Pten^PEΔ/Δ^*prostates. a-b) Genotypes and expression levels are color coded. c) Expression levels (log2, FPKM) of *Ccl3, Ccl8* and *Il1b* are significantly (*Ccl3* p=2.4e-04; *Ccl8*, p=9.7e-05 and *Il1b*, p=5.0e-03) reduced in EpCAM^+^ cells of *Jun^PEΔ/Δ^;Pten^PEΔ/Δ^*prostates. Significance was determined by an unpaired two-sided t-test. d) Single-sample GSEA analysis using the M5 signature of Broad Institute’s molecular signature database (MsigDB) revealing enrichment of macrophage- and neutrophil-associated properties in *Pten^PEΔ/Δ^* compared to *Jun^PEΔ/Δ^;Pten^PEΔ/Δ^* prostates. e) Western blot analysis of NLRP3 and non-cleaved Pro-IL-1β in all four experimental groups in biological replicates. β-ACTIN served as loading control. f) Multiplex immunoassay of homogenized prostate samples of 19-week-old *wt*, *Jun^PEΔ/Δ^, Pten^PE^*^Δ/Δ^ and *Jun^PEΔ/Δ^;Pten^PEΔ/Δ^*animals for analysis of IL-1β levels in pico grams (pg)/ml of indicated biological replicates. Statistical testing was done with one-way Anova, significant p-values are indicated.

Cells of the innate immune system, including neutrophil granulocytes, mast cells and macrophages serve as the primary defense against infections and consequently recruit T and B cells to infection sites [46]. Among the differentially expressed genes (DEGs) of *Pten^PE^*^Δ/Δ^ versus *Jun*^PEΔ/Δ^; *Pten*^PEΔ/Δ^ prostates, we identified neutrophil movement-specific gene signatures that play a crucial role in the recruitment of immune cells (Fig. 4b) [47]. We observed that cytokines involved in chemotaxis of immune cells such as *Ccl3*, *Ccl8* and *Il1b* were significantly deregulated between groups (Fig. 4c). To further dissect the potentially involved immune cell subsets, we conducted single sample GSEA using the M5 ontology gene sets signature from the molecular signature database (MsigDB). We identified enrichment of macrophage- and neutrophil-specific gene signatures characterized by cellular activities such as migration, activation/differentiation and enhanced expression indicating production of MIP1α/CCL3 and GM-CSF. Moreover, single sample GSEA revealed processes related to other immune cell subsets such as mast cells, myeloid cells and CD8^+^ T cells that were significantly enriched in *Pten^PEΔ/Δ^* compared to *wt* prostates and depleted in *Jun*^PEΔ/Δ^; *Pten*^PEΔ/Δ^ (Fig. 4d). This implicates JUN in the control of inflammatory states during PCa progression. We validated the JUN-dependent regulation of IL-1β and NLRP3 both involved in the regulation of inflammatory response processes [48] by immunoblot and cytokine analyses (Figs. 4e-f).

To further examine the apparent shifts in immune system-related transcriptomic signatures, we assessed granulocytic or lymphocytic cell infiltrations based on microscopic characteristics in H&E staining of all four genotypes (Supplementary Fig. 4b). We detected no or low-grade infiltration by inflammatory cells in *wt* and *Jun^PE^*^Δ/Δ^ specimens. In contrast, *Pten^PE^*^Δ/Δ^ mouse prostates exhibited increased levels of high- and middle-grade infiltrations, which were significantly mitigated in *Jun*^PEΔ/Δ^; *Pten*^PEΔ/Δ^ prostates. Increased immune cell infiltration of *Pten^PE^*^Δ/Δ^ prostates as identified by histo-pathological analysis therefore supported the results of transcriptome profiling. This highlights the importance of JUN in the regulation of senescence-associated inflammation in *Pten*-deficient PCa.

### Epithelial JUN deficiency modulates the migration of innate immune cells from the periphery

To investigate the distribution and abundance of infiltrating immune cells, we performed IHC stainings. Neutrophils and inflammatory monocytes were stained using the antibody clone NIMP-R14, which targets the specific cell surface markers and differentiation antigens Ly-6G and Ly-6C (Fig. 5a). In *Pten^PE^*^Δ/Δ^ prostates, we observed high numbers of neutrophils migrating from the blood vessels across the stroma into the epithelium, where they predominantly accumulated, and subsequently advanced into the lumen. In *Jun*^PEΔ/Δ^; *Pten*^PEΔ/Δ^ prostates, we detected significantly (p<1.0e-04) less neutrophils in the stroma and epithelium, but the migration patterns remained consistent with *Pten^PE^*^Δ/Δ^ tumors (Fig. 5b). In contrast, macrophages, stained by the marker F4/80 were primarily located in the stroma, with no significant differences between the groups (Figs. 5c-d). We observed significantly (p=4.0e-04) less macrophages infiltrating the epithelium in prostates with additional deficiency of *Jun*. In conclusion, *Pten^PE^*^Δ/Δ^ displayed a highly immune infiltrated phenotype, which was substantially reverted in prostates with additional deficiency of *Jun*. This observation suggests that JUN may be essential for tumor cell recognition by innate and consequently adaptive immune cells.

**Figure 5:**
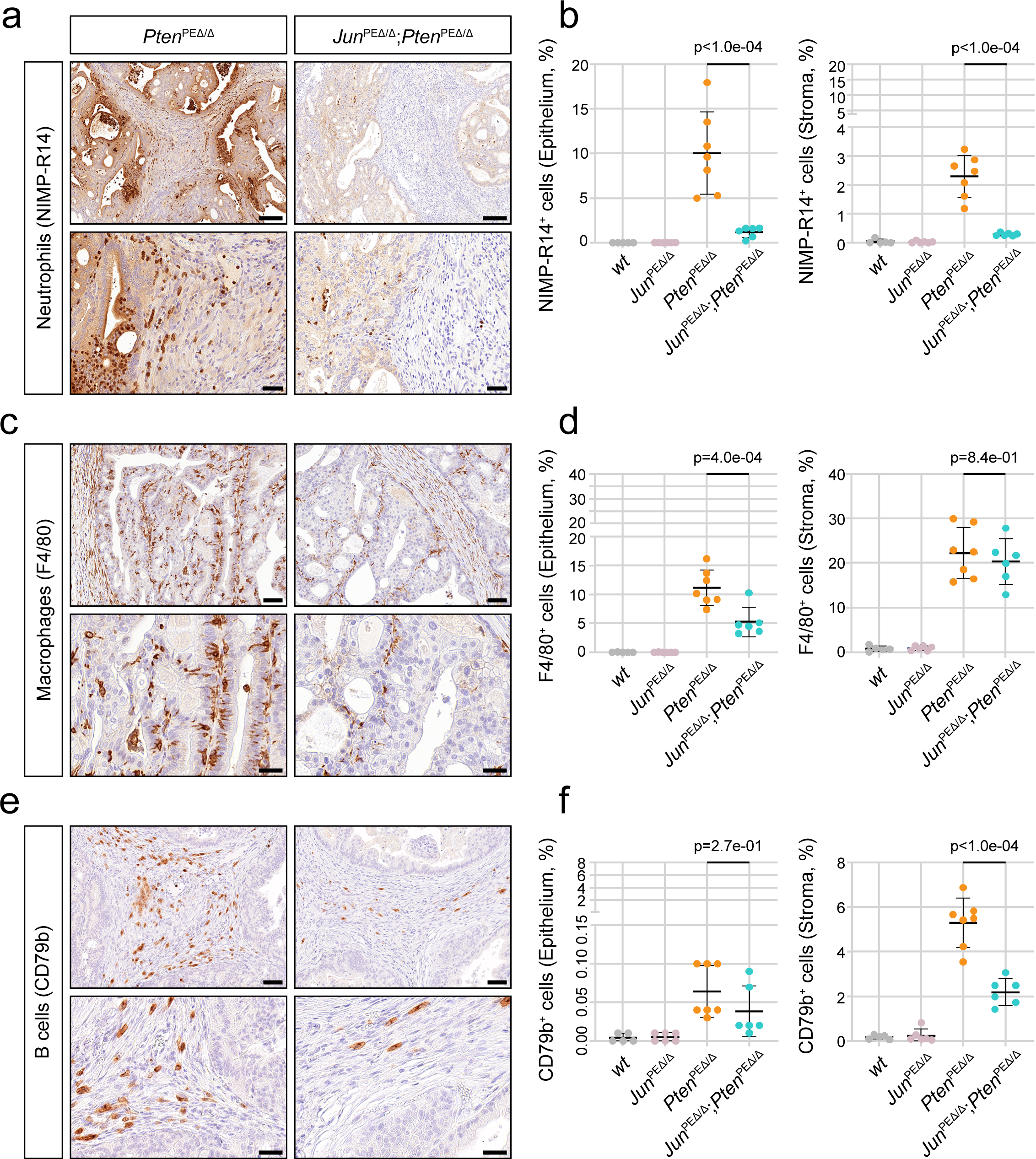
Histological analysis of infiltrating immune cells reveals downregulated innate immune response in *Jun^PEΔ/Δ^;Pten^PEΔ/Δ^* prostates. a) Representative images of IHC stainings of NIMP-R14, a pan-marker of neutrophil granulocytes, indicating high neutrophil infiltration of *Pten^PEΔ/Δ^* prostates, reverted by the additional loss of Jun in *Jun^PEΔ/Δ^;Pten^PEΔ/Δ^*prostates. Top row: 20.0x magnification, scale bar represents 150 µm; Bottom row: 63.0x magnification, scale bar represents 40 µm. b) Quantification of NIMP-R14^+^ neutrophils in epithelium (left) and stroma (right). A significantly decreased (p<1e-04) infiltration of neutrophils in tumors and adjacent stroma of *Jun^PEΔ/Δ^;Pten^PEΔ/Δ^* prostates is evident. c) Representative images of IHC stainings for the pan-marker of macrophages F4/80. A high infiltration of *Pten^PEΔ/Δ^* prostates and adjacent stroma by macrophages is evident and reverted by the additional loss of Jun in *Jun^PEΔ/Δ^;Pten^PEΔ/Δ^* prostates. Top row: 40.0x magnification, scale bar represents 60 µm; Bottom row: 100.0x magnification, scale bar represents 30 µm. d) Quantification of F4/80^+^ macrophages in epithelium (left) and stroma (right). A significantly decreased (p=4e-04) infiltration of macrophages in tumors but not adjacent stroma (p=8.3e-01) of *Jun^PEΔ/Δ^;Pten^PEΔ/Δ^* prostates is evident. e) Representative images of IHC stainings of B cell infiltration using the pan-marker CD79b. A high infiltration of stroma adjacent to *Pten^PEΔ/Δ^* prostates by CD79b^+^ B cells is evident and reverted by the additional loss of *Jun* in *Jun^PEΔ/Δ^;Pten^PEΔ/Δ^*prostates. Top row: 40.0x magnification, scale bar represents 60 µm; Bottom row: 100.0x magnification, scale bar represents 30 µm. f) Quantification of B cells in epithelium (left) and stroma (right). B cell infiltration as observed in the stroma of *Pten^PEΔ/Δ^* prostates was significantly decreased (p<1e-04) in *Jun^PEΔ/Δ^;Pten^PEΔ/Δ^*prostates. b, d, f) Statistical significance between *Pten^PEΔ/Δ^* and *Jun^PEΔ/Δ^;Pten^PEΔ/Δ^* groups are indicated.

Neutrophils attract T cells to the site of inflammation via secretion of chemokines such as CCL2 and CCL5 [49,50]. We utilized multiplex IHC to discern the T cell subsets, employing a marker panel consisting of CK/CD3/CD4/CD8/CD45/PD-1/DAPI. We observed various T cell subpopulations (T helper (CD4^+^) and cytotoxic T cells (CD8^+^), PD-1 positive and negative) mainly in the stroma and to a lesser degree in the epithelium. We did not observe a significant effect of *Jun* deficiency on any of the subpopulations (Supplementary Figs. 5a-b and data not shown). Additionally, we investigated the infiltration of B cells, stained by CD79b. B cells were found almost exclusively in the stroma, with significantly less infiltration in *Jun*^PEΔ/Δ^; *Pten*^PEΔ/Δ^ compared to *Pten^PE^*^Δ/Δ^ prostates (Figs. 5e-f). In summary, IHC validated the JUN-dependent modulation of the immune cell compartment, particularly affecting innate immune cells. This phenotype was likely provoked by a JUN-dependent regulation of neutrophil attracting chemokines such as IL-1β.

### Increased expression of SASP factors is correlated with prolonged survival in prostate cancer

To translate our findings to the human disease, we investigated a potential correlation between JUN and SASP factors and a dependence of immune cell infiltration on AP-1 factors using a human PCa dataset [40]. We observed a significant positive correlation of *JUN* and levels of *IL1B* (R=0.47, p<2.2e-16), *CCL8* (R=0.42, p=2.2e-16) and *CCL3* (R=0.51, p<2.2e-16) as well as a weak but significant correlation of *JUN* and *F4/80* (*ADGRE1*, R=0.20, p=2.0e-16) (Figs. 6a-d). Next, we assessed whether AP-1 factors may directly determine the degree of immune cell infiltration. We used the ESTIMATE tool that allows calculation of a score to predict immune cell infiltration based on expression levels of specific signature genes [51]. We ranked cancer genome atlas prostate adenocarcinoma (TCGA-PRAD) [40] samples based on their ImmuneScores, distinguishing tumors with high and low infiltration levels and examined genes altered in our murine model. We observed that levels of *JUNB* and *FOSL1* but not *JUN*, *JUND*, *FOS* or *FOSB* were associated with a high ImmuneScore, potentially indicating a prognostic relevance of these markers (Supplementary Fig. 6a). We also detected considerable correlation of the SASP factors *IL1B* (R=0.52, p<2.2e-16), *CCL8* (R=0.45, p<2.2e-16), *CCL3* (R=0.45, p<2.2e-16) and *ADGRE1* (R=0.67, p<2.2e-16) with ImmuneScore (Figs. 6e-h). A correlation analysis validated the relationship of all AP-1 factors investigated with immune relevant SASP markers, particularly *IL1B*, *CCL3*, *CCL8* but not *IL-18* (Supplementary Fig. 6b). Assuming that JUN mediates tumor-suppressor activity via positive regulation of SASP factors required for the recruitment of immune cells, we expected that high levels of SASP factors may be associated with favorable prognosis. As observed for *JUN,* high expression of *IL1B* [HR=0.41 (0.22 − 0.78), logrank p=4.5e-03], *CCL8* [HR=0.53 (0.3 − 0.96), logrank p=3.4e-02] and *CCL3* [HR=0.47 (0.25 − 0.89), logrank p=1.8e-02] indeed correlated with reduced BCR and significantly improved patients survival [40] (Figs. 6i-k). Finally, we investigated a potential relationship between JUN and STAT3 activation and explored STAT3’s role in immune modulation. We correlated reverse-phase protein array (RPPA) [40] of pSTAT3^Y705^ with levels of JUN and IL-1β. We observed a significant correlation of JUN (R=0.47, p<2.2e-16) and IL-1β (R=0.48, p=2.2e-16) with pSTAT3^Y705^ (Supplementary Fig. 6c). As observed for JUN, PCa exhibiting a high (>7) Gleason score showed reduced levels of pSTAT3^Y705^ (p=1.6e-02) when compared to low risk tumors (Gleason score ≤7) (Supplementary Fig. 6d). These results might hint at concerted mechanisms of both transcriptional regulators. In summary, we propose that levels of JUN determine progression stages of prostate tumors by modulating the immune response through regulation of cytokines and interleukins analogous to our results from the *Jun*-deficient murine PCa model.

**Figure 6:**
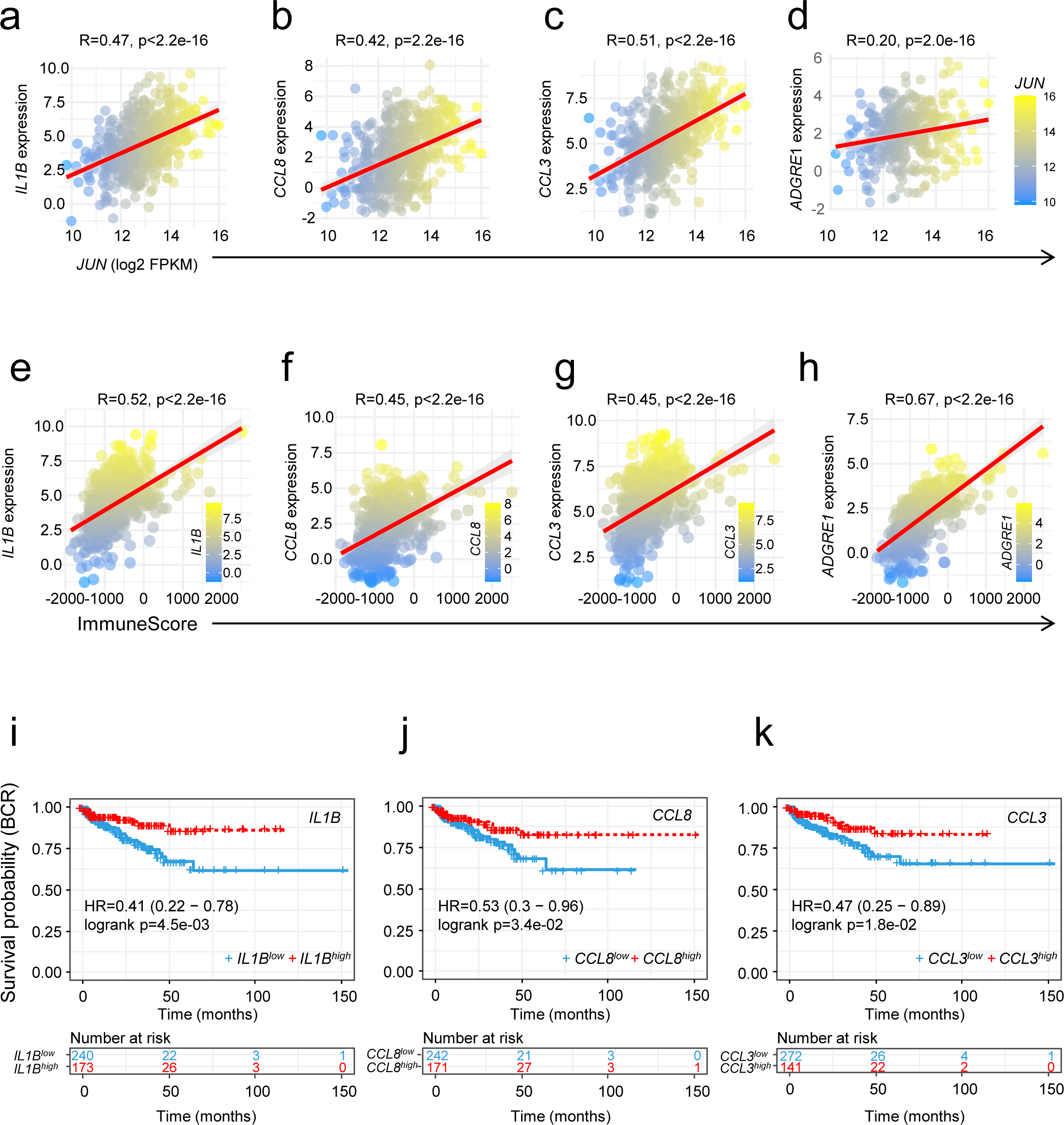
Expression of immune cell-attracting chemokines *CCL3* and *CCL8* correlates with levels of *JUN* in patient datasets. Correlation of *JUN* expression and levels of *IL1B* (a), *CCL8* (b), *CCL3* (c) and pan macrophage marker *ADGRE1/F4/80* (d). The *JUN* level (log2 FPKM) is color coded (a-d). Correlation analysis of IL-1β (e), CCL8 (f), CCL3 (g) and ADGRE1/F4/80 (h) to the ImmuneScore in human PCa datasets (TCGA-PRAD [40]). The level of correlation is color coded (e-h). Kaplan-Meier survival analyses of TCGA-PRAD tumors [40] showing that high expression of IL-1β (i), CCL8 (j) and CCL3 (k) correlated with favorable survival and reduced probability of BCR.

## Discussion

PCa is among the most frequently diagnosed malignancies in men worldwide and a significant number of patients progress to advanced and lethal stages. The mortality linked to metastatic PCa highlights the pressing need to elucidate its intricate mechanisms and pinpoint viable therapeutic interventions. Despite this urgency, the cellular mechanisms and environmental contexts that control PCa development and progression remain incompletely understood. Loss of *PTEN* is evident in 20% of primary human prostate carcinomas and escalates in 50% of metastatic CRPC [4]. Comparable to the human situation, *Pten* loss leads to the formation of precancerous lesions in PE cells in mouse models [52,53]. Aggressive carcinomas develop only in the presence of additional mutations [54], such as abnormal expression of ERG [55], loss of IL-6/STAT3 functionality [12,13], dysfunction of the methyltransferase Kmt2c [28] or activation of the RAS/MAPK cascade [54,56]. While several studies indicate that augmented JUN expression drives PCa progression [14,57], the functional role of JUN and AP-1 TFs in PCa remains controversial. Intriguingly, genetic disruption of *JUNB* and *FOS in vivo* accelerated the progressive phenotype of *PTEN*-deficient PCa [22,58]. This suggests a context dependent tumor-suppressive role, rather than a driving function of AP-1 TFs in prostate cancer progression [59].

In the present study, our focus was to delineate the role of JUN in PCa. We first examined JUN levels in clinical PCa samples and analyzed *JUN* patterns across varying progression stages from three publicly available datasets [39,41,42]. We found that *JUN* expression increased in tumors relative to normal prostates but the levels of *JUN*, *FOS* and *JUNB* significantly decreased with progression of PCa. This suggests that high JUN levels may protect from development of progressive disease, a hypothesis further supported by the survival rates of patients harboring high *JUN* expressing tumors. Encouraged by these findings, we studied the functional role of JUN in a murine PCa model, characterized by homozygous loss of *Pten* (*Pten^PEΔ/Δ^*) [26,43]. Mirroring the early prostatic intraepithelial neoplasia (PIN) stages of human PCa, JUN was significantly upregulated in *Pten^PEΔ/Δ^* prostates. While depletion of *Jun* alone had no effect on the morphological architecture and growth of the prostate, epithelial cells of *Pten^PEΔ/Δ^* prostates developed hyperplasia, subsequently formed prostate adenocarcinoma and rapidly progressed upon additional deletion of *Jun*. The aggressive phenotype observed in *Jun^PEΔ/Δ^*;*Pten^PEΔ/Δ^*prostates resulted in decreased survival of mice and increased prostate weight and size. We did not detect signs of severe organ dysfunction, systemic inflammation or metastatic disease (data not shown). The TME is a dynamic system characterized by chronic inflammation and participation of diverse host components, but plays a pivotal role in cancer progression [60]. Within the TME, immune cells such as tumor-associated macrophages (TAM) and tumor-associated neutrophils (TAN) both foster cancer progression or combat tumor cells, underscoring their dual roles in tumorigenesis [61–63]. Central to this environment is the SASP, where senescent cells release a plethora of inflammatory mediators. SASP-driven effects often culminate in the immune-mediated clearance of potential tumorigenic cells, a process termed “senescence surveillance” [64,65]. Our histopathologic examination of *Pten^PEΔ/Δ^*PCa samples revealed significant enrichment of neutrophils and macrophages that infiltrated the tumors and adjacent stroma. Concurrent deletion of *Jun* strikingly reduced tumor infiltration with neutrophils and macrophages and accelerated tumor growth. Transcriptomic analyses of *Pten^PEΔ/Δ^* and *Jun^PEΔ/Δ^*; *Pten^PEΔ/Δ^*prostates revealed a JUN-dependent modulation of SASP-associated genes, but we did not identify compensatory upregulation of other AP-1 members as it has been described upon inactivation of FOS [58]. Recent findings underscore AP-1 TF’s pioneering role at genomic enhancers. Binding of AP-1 and in particular of JUN, imprints the senescence enhancer landscape to regulate factors that are needed for the execution of senescence-controlling programs [23]. In line with these results, we propose that loss of *Pten* coupled with an increase in JUN levels likely instigates a JUN-driven SASP phenotype which is accompanied by widespread changes in gene expression. SASP involves the expression and secretion of inflammatory cytokines such as CCL3, CCL8, IL-1β and growth factors [28,66] which subsequently recruit immune cells such as neutrophils, macrophages and T cells [10,67,68]. Consequently, *Jun* depletion in *Pten*-deficient prostates may abrogate SASP and impede recruitment of neutrophils and macrophages as well as tumor cell clearance by macrophages and dendritic cells [9,64]. We thus propose JUN as a key regulator of SASP which is in line with a previous study [23]. Their observations linked JUN depletion to diminished inflammatory responses which reverted the senescent/SASP phenotype of RAS-OIS fibroblasts to a proliferating phenotype [23]. Furthermore, GM-CSF, a direct JUN target, has been shown to amplify macrophage and neutrophil immune responses [69] and modulate pro-inflammatory cytokine secretion such as TNFα and IL-6 [70].

Another intriguing mechanism of JUN-dependent modulation of the immune phenotype in PCa may depend on STAT3 levels. Our previous work identified activation of STAT3 and a p19^ARF^– MDM2–p53 axis to induce senescence upon *Pten* depletion [12]. Consistently, *Jun* loss was associated with decreased IL-6-JAK-STAT3 signaling, evidenced by significantly reduced pSTAT^Y705^ levels in *Jun^PEΔ/Δ^*; *Pten^PEΔ/Δ^* prostates. ENCODE database exploration [71] revealed mutual promoter binding sites for JUN and STAT3 suggesting a potential JUN-STAT3 interplay in guiding senescence pathways in PCa (data not shown). This interplay is supported by results of a STAT3 binding analysis in CD4^+^ T cells, which suggests that STAT3 directly regulates the expression of *Jun* and *Fos* and may potentially function in a positive feedback loop [72]. Therefore, therapeutic activation of STAT3 potentially causes SASP factor modulation and may elevate JUN levels in tumors, thereby restricting tumor progression and enhancing PCa patient survival.

## Conclusions

In summary, our data suggest that JUN functions as a pivotal regulator of SASP in PCa, orchestrating the recruitment dynamics of TAMs and TANs within the TME. Given the indispensable role of robust SASP in immune surveillance of preneoplastic anomalies, its therapeutic modulation presents intricate challenges. Our recent investigations have shown the potential of the antidiabetic agent metformin, which curtails multiple pro-inflammatory SASP components by inhibiting NF-κB nuclear translocation [73]. Metformin increases STAT3 in advanced PCa cases, leading to significant tumor growth attenuation, underscored by reduced mTORC1/CREB and AR levels in a PCa murine model [13]. The interplay between JUN and STAT3 might represent a key mechanism that could be exploited for therapeutic advances.

## List of Abbreviations

AP-1: Activator protein-1
AR: Androgen receptor
BCR: Biochemical recurrence
CRPC: Castration resistant prostate cancer
DEG: Differentially expressed gene
EpCAM: Epithelial cell adhesion molecule
FDR: False discovery rate
FFPE: Formalin-fixed paraffin embedded
GO: Gene ontology
GSEA: Gene set enrichment analysis
H&E: Hematoxylin and eosin
HR: Hazard ratio
IHC: Immunohistochemistry
IF: Immunofluorescence
JNK: JUN N-terminal kinase
MsigDB: molecular signature database
NEPC: Neuroendocrine prostate cancer
NES: Normalized enrichment score
OIS: Oncogene induced senescence
PIN: Prostatic intraepithelial neoplasia
PICS: PTEN-loss induced cellular senescence
PbCre: Probasin Cre
PCa: Prostate cancer
PCA: Principal component analysis
PE: Prostate epithelium
PI3K: Phosphoinositide 3-kinase
PTEN: Phosphate and Tensin Homologue
RFS: Relapse free survival
RNA-seq: RNA sequencing
RPPA: Reverse-phase protein array
SASP: Senescence-associated secretory phenotype
STAT3: signal transducer and activator of transcription
3 TAM: Tumor-associated macrophage
TAN: Tumor-associated neutrophil
TCGA-PRAD: Cancer Genome Atlas Prostate Adenocarcinoma
TF: transcription factor
TMA: Tissue microarray
TME: Tumor microenvironment

## Declarations

### Ethics approval and consent to participate

Institutional Review Board Statement: The use of clinical material was approved by the Research Ethics Committee of the Medical University Vienna, Austria (1877/2016) and conducted in adherence to the Declaration of Helsinki protocols. Patient consent was waived due to the completely anonymized, retrospective nature of the study.

All animal studies were reviewed and approved by the Federal Ministry Republic of Austria for Education, Science and Research and conducted according to regulatory standards (BMWFW-66.009/0144-WF/II/3b/2014 and the amendments BMWFW-66.009/0063-WF/V/3b/2017, BMWFW-66.009/0137-WF/V/3b/2019, BMWFW-66.009/0359-V/3b/2019, 2020-0.016.881, 2020-0.659.052 and 2022-0.325.014).

### Consent for publication

Not applicable.

### Availability of data and materials

The RNA-seq dataset supporting the conclusions of this article is available in the GEO repository with the accession number GSE242433, and is publicly available as of date of publication. The following publicly available datasets were used: GSE21034 [39], TCGA-PRAD [40], GSE6919 [41], E-MTAB-9930 [42]. Expression levels of AP-1 factors of PCa subtypes were retrieved from a minimum dataset (vst-normalized expression data), (Zenodo repository) [42].

### Authors contributions

Conceptualization: TR, MR, CS, SL, LK. Formal Analysis: TR, KT, MO, MB, HAN, VB. Funding Acquisition: LK, SP. Investigation: TR, MR, CS, RZ, CP, DL, AA, TL, PK, SH, MS, SS, MO, MB, HAN, IG, OM, SL. Methodology: MR, CS, RZ, CP, AA, TL, SMi, MT, NSH, BT, VB, BS, GE, SL. Project administration: TR, MR, CS, SL, LK. Resources: TR, MR, CS, RM, JT, SMa, FA, SP, JP, GE, SL, LK. Supervision: SL, LK. Validation: TR, MR, CS, KT, PK, SH, MS, MO, FS, BT, VB, OM, GE. Visualization: TR, MR, CS, RZ, CP, AA, MB, HAN, TL. Writing – original draft: SL, TR, MR. Writing – review & editing: TR, MR, CS, TL, SS, RM, SMa, FA, BS, OM, GE, SL, LK. All authors read and approved the final manuscript.

## Acknowledgements

This research was supported using resources of the VetCore Facilities (VetImaging, Genomics and Transcriptomics) of the University of Veterinary Medicine Vienna. We acknowledge the Core Facility Genomics and Core Facility Bioinformatics supported by the NCMG research infrastructure (LM2023067 funded by MEYS CR) for their support with the bioinformatic analysis of scientific data presented in this paper. We would like to thank Anton Jäger, Medical University of Vienna for macroscopic image generation and support.

## Authors’ information

Torben Redmer, Martin Raigel and Christina Sternberg contributed equally to this work as first authors. Sabine Lagger and Lukas Kenner contributed equally to this work as last authors.

---

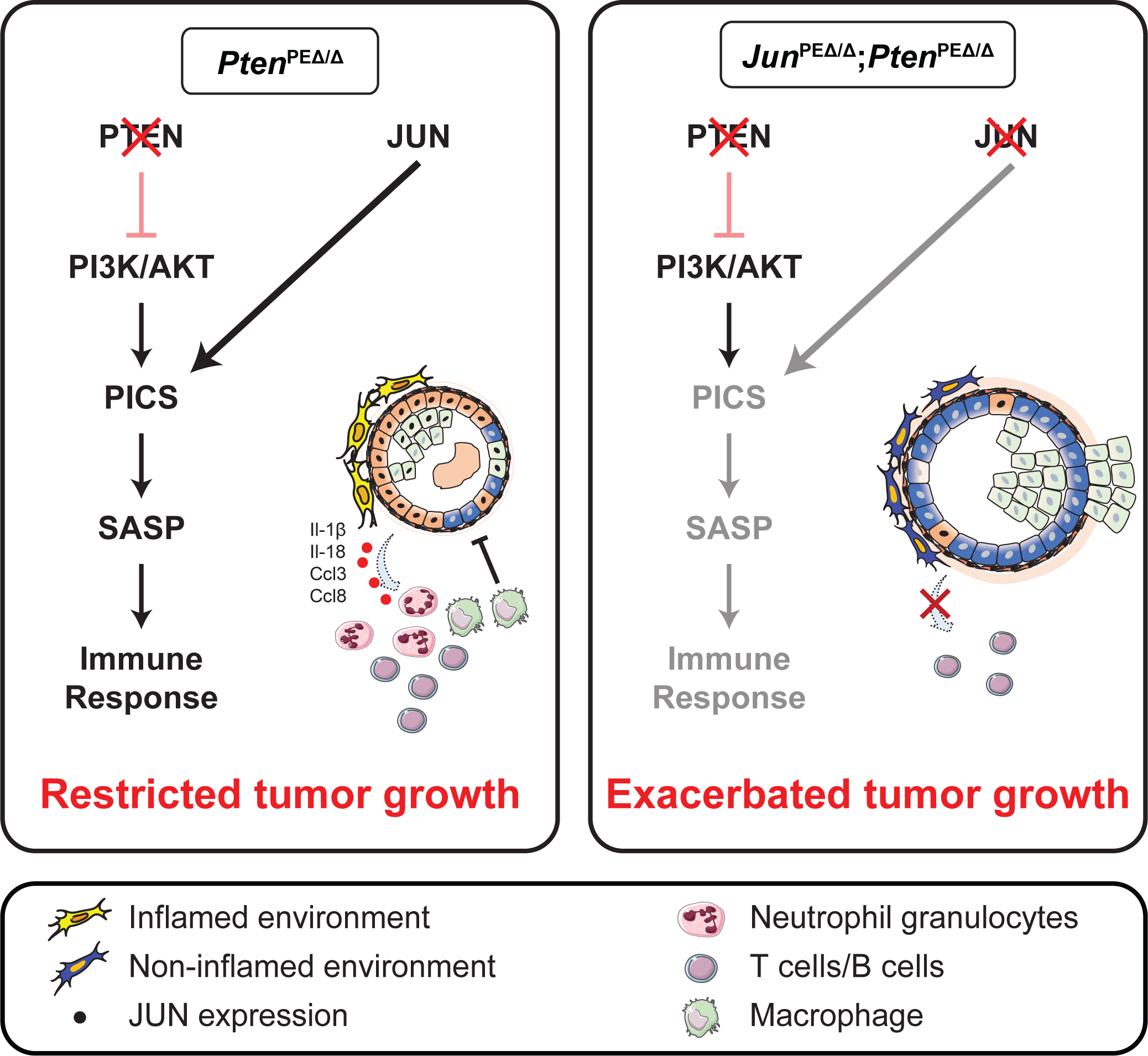

