## Supplementary Figures for "JUN mediates senescence and immune cell recruitment to prevent prostate cancer progression"

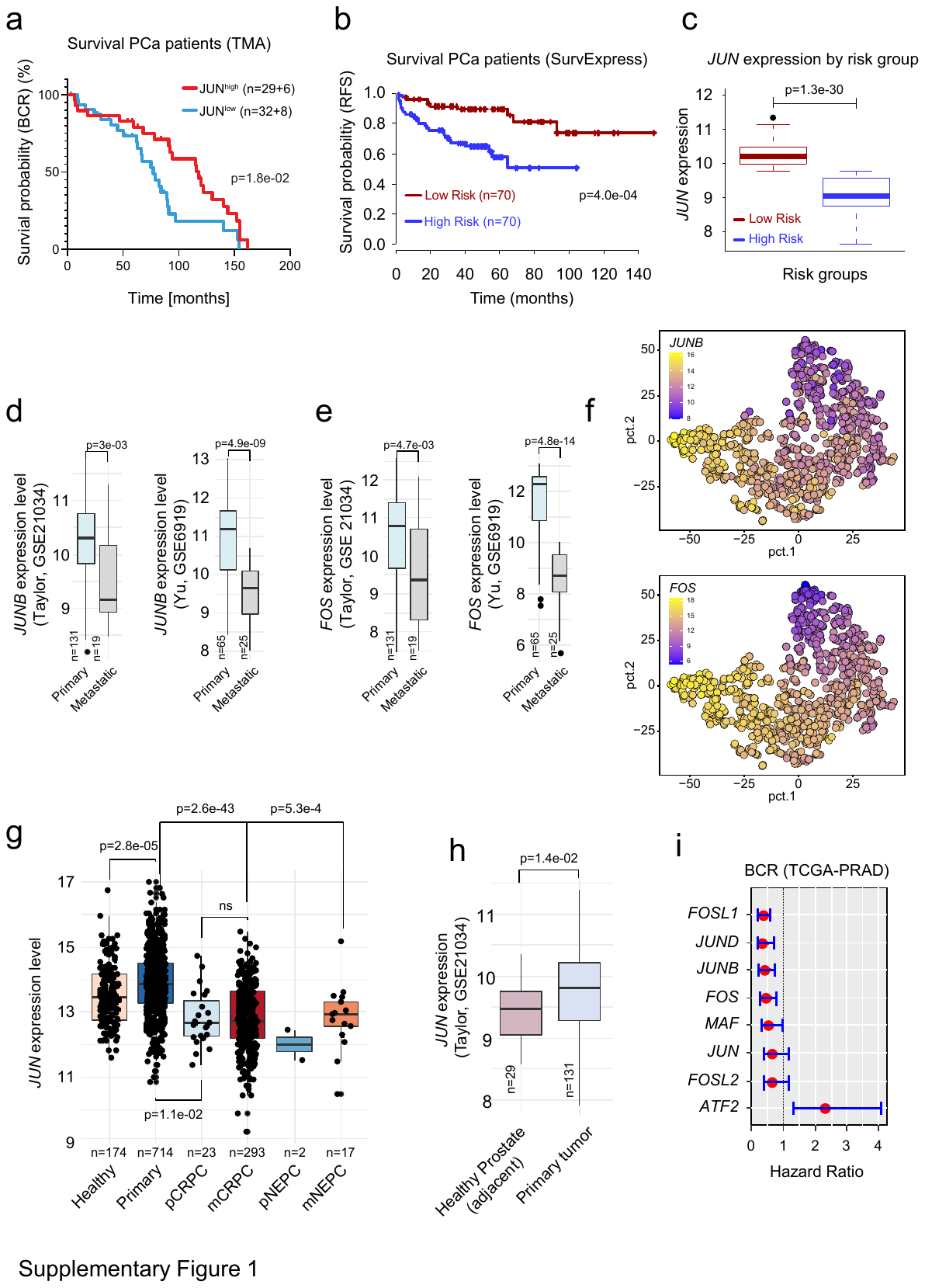

**Supplementary Figure 1: Levels of JUN define progression stages in prostate cancer.** a) Kaplan-Meier survival analysis of human prostate tumors (n=75) used for analysis of TMAs stained with an antibody against JUN. Patient groups stratified by presence or absence of JUN show significantly (p=1.8e-02) different BCR-free survival in (%). b) Survival curves (RFS) of PCa patient RNA-seq data [39] were analysed with the SurvExpress web tool [32]. The groups were stratified according to the prognostic index (PI) into high (red) and low (blue) risk groups. The group comparison was performed with a logrank test (p=4.0e-04). c) *JUN* mRNA expression was surveyed in the individual patient risk groups generated in b). d-e) *JUNB* and *FOS* mRNA levels in prostate tumors comprised in the Taylor [39] (p=3.0e-03 and p=4.7e-03) and Yu datasets [41] significantly (p=4.9e-09 and p=4.8e-14) discriminated primary and metastatic tumors. f) Overlay of *FOS* and *JUNB* mRNA expression with the principal component analysis (PCA) from Fig. 1f. *FOS* and *JUNB* levels are color coded from high expression (yellow) to low expression (blue). g) Investigation of *JUN* mRNA expression in normal prostate tissue, primary adenocarcinoma and primary (p) and metastatic (m) CRPC and NEPC. Data were retrieved from [42]. h) *JUN* levels significantly (p=1.4e-02) discriminate healthy adjacent prostate tissue (n=29) and primary (n=131) prostate tumors. Data were retrieved from [39]. i) Representation of hazard ratios (95% confidence interval) of AP-1 family genes determined by Kaplan-Meier analyses with the KMplot tool. The TCGA-PRAD data were used for the analysis [40]. Hazard ratios refer to BCR. In d, e, g, h statistical significance was determined by an unpaired, two-sided t-test.

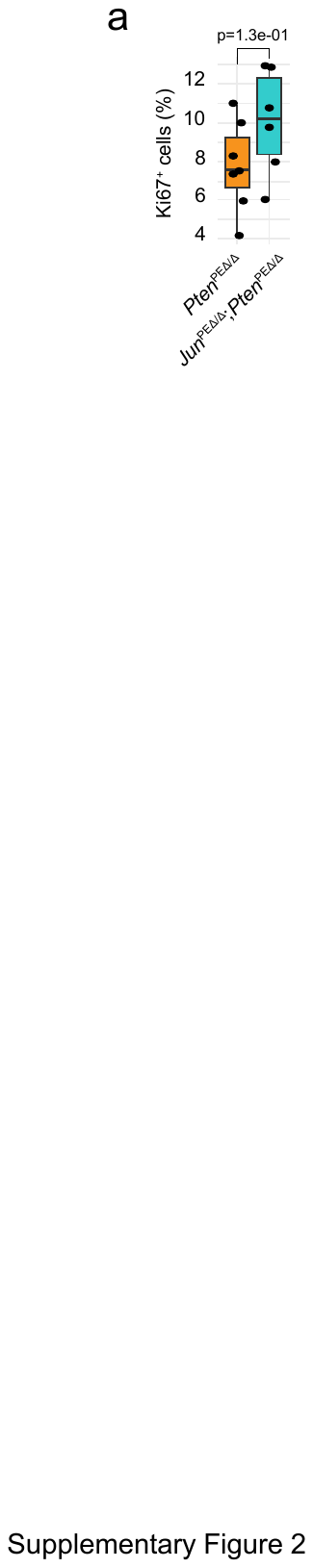

**Supplementary Figure 2: JUN does not control cellular proliferation in *Pten*-deficient prostate tumors** a) Box plot depicting the percentage of Ki67^+^ epithelial cells assessed from IHC staining of *Pten^PEΔ/Δ^* (n=7) and *Jun^PEΔ/Δ^;Pten^PEΔ/Δ^* (n=6) prostate samples. Data points represent the mean of four individual regions of interest with a radius of 150 µm analyzed by manually counted positive epithelial cells.

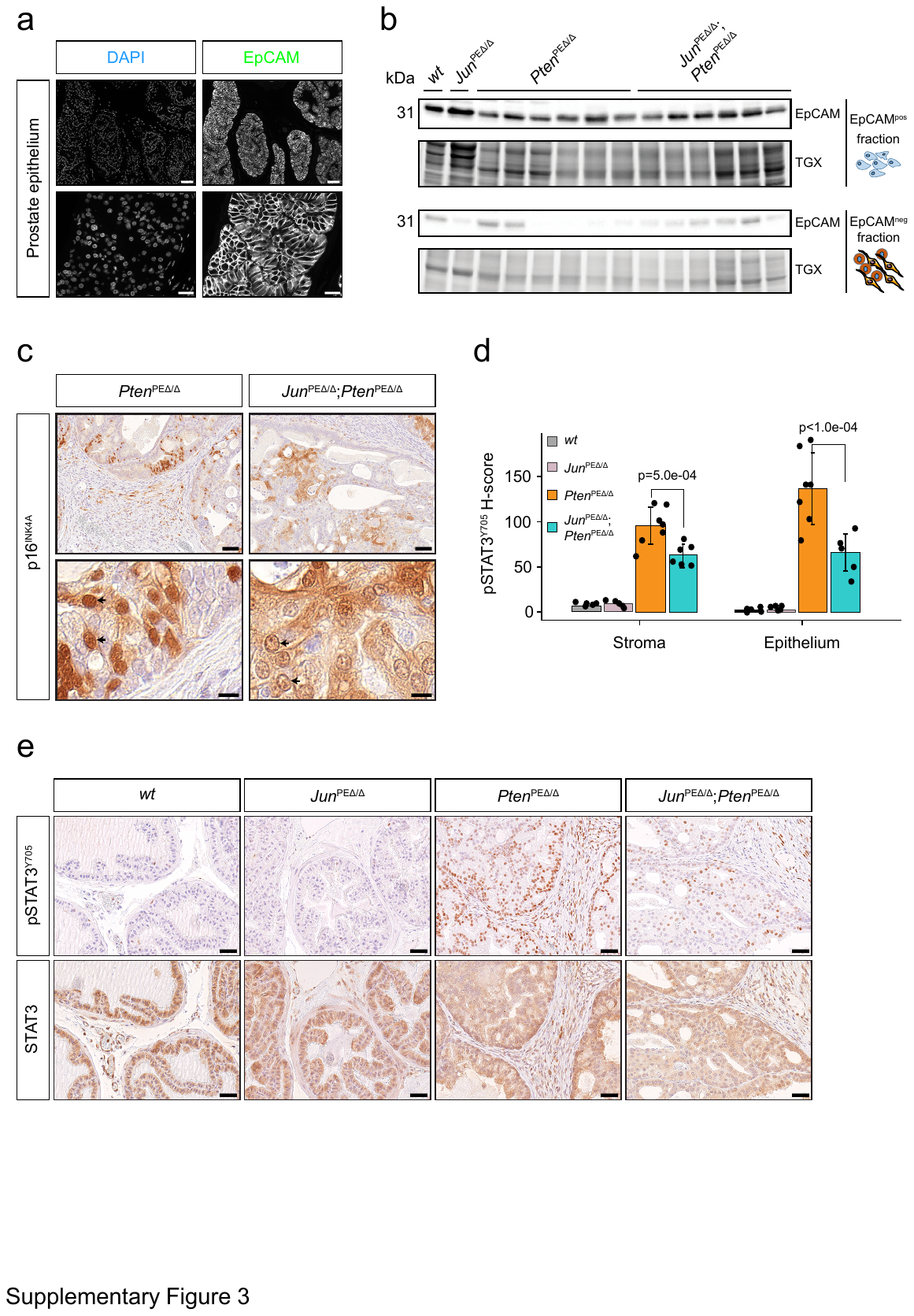

**Supplementary Figure 3: EpCAM expression separates epithelial and stromal cells.** a) Single channel grayscale representative images of DAPI/EpCAM immunofluorescence (IF). Top row: 40.0x magnification, scale bar represents 60 µm; Bottom row: 147.5 x magnification, scale bar represents 20 µm. b) Immunoblot analysis with an antibody against EpCAM to survey separation of mouse prostate sample single cell suspensions in EpCAM positive and negative fractions after magnetic cell sorting in all experimental groups. TGX stain free technology is used as loading control. c) Representative pictures of IHC stainings with antibodies against p16^INK4A^ in *Pten^PE^*^Δ/Δ^  and *Jun^PEΔ/Δ^;Pten^PEΔ/Δ^* animals. Top row: 40.0x magnification, scale bar represents 60 µm; Bottom row: 300.0 x magnification, scale bar represents 10 µm. d) Quantification of IHC staining with an antibody against phosphorylated STAT3 at tyrosine Y705 (pSTAT3^Y705^) in indicated biological replicates of 19-week-old *wt*, *Pten^PE^*^Δ/Δ^, *Jun^PEΔ/Δ^* and *Jun^PEΔ/Δ^;Pten^PEΔ/Δ^* animals according to H-score. Stroma and epithelium were analysed individually by digital pathology software on whole slide scans. Statistical testing was done with one-way Anova with Turkey’s multiple comparison and statistical significance between *Pten^PE^*^Δ/Δ^ and *Jun^PEΔ/Δ^;Pten^PEΔ/Δ^* animals is shown. e) Representative pictures of IHC stainings with antibodies against pSTAT3^Y705^ (upper panels) and total STAT3 (lower panels) in all four experimental genotypes. Images were taken in a 63.0x magnification and the scale bar represents 40 µm.

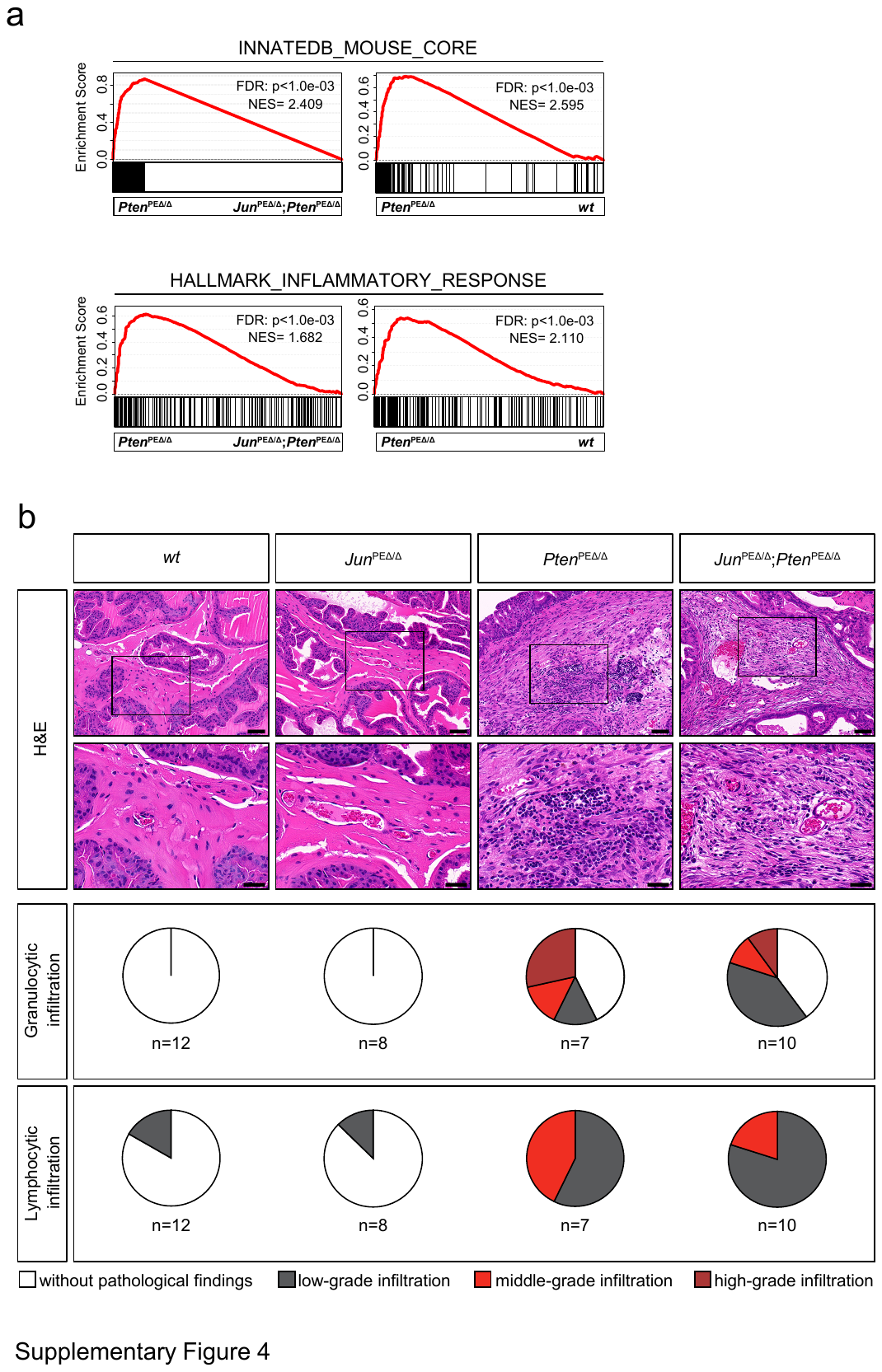

**Supplementary Figure 4: JUN level determines stages of immune cell infiltration.** a) GSEA enrichment analysis using InnateDB_mouse_core (upper panels) and HALLMARK_INFLAMMATORY_RESPONSE gene sets in *Pten^PE^*^Δ/Δ^ versus *Jun^PEΔ/Δ^;Pten^PEΔ/Δ^* and *Pten^PE^*^Δ/Δ^ versus *wt* animals. b) Upper panels: Representative H&E images depicting the morphology of prostates and infiltration of granulocytes and lymphocytes in 19-week-old animals. Top row: 40.0x magnification, scale bar represents 60 µm; Bottom row: 100.0 x magnification, scale bar represents 30 µm. Lower panels: The sections were evaluated by an independent pathologist and the level of immune cell infiltration (white = without infiltration; grey = low-grade; red = middle-grade; dark red = high-grade infiltration) of *wt* (n=12), *Jun^PEΔ/Δ^* (n=8), *Pten^PEΔ/Δ^* (n=7) and *Jun^PEΔ/Δ^;Pten^PEΔ/Δ^* (n=10) prostates were assessed and summarized in pie charts.

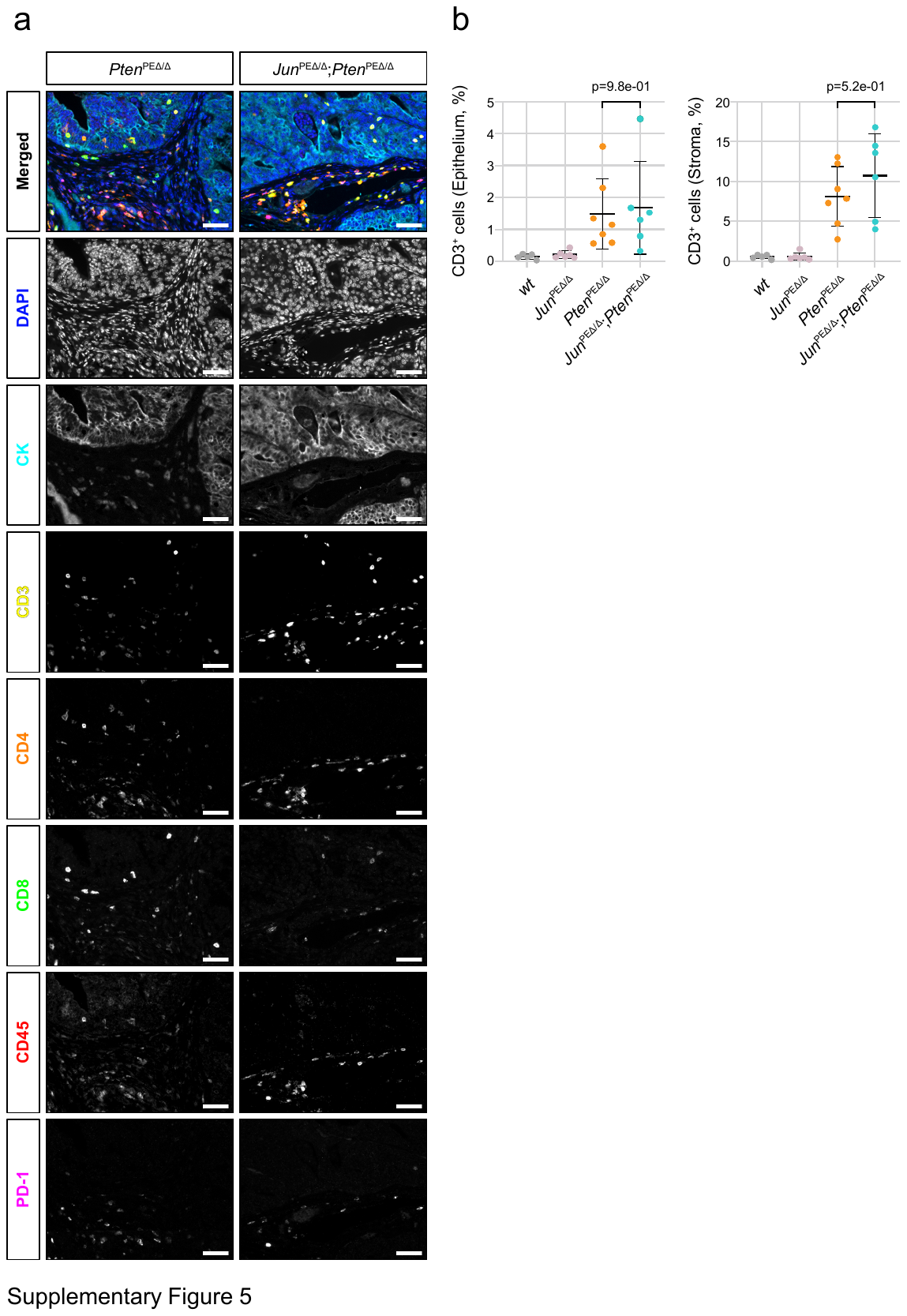

**Supplementary Figure 5: The composition of T cell subsets is not affected by *Jun*-deficiency.** a) Merged and single channel grayscale representative images of multiplex IHC of sections of *Pten^PEΔ/Δ^* and *Jun^PEΔ/Δ^;Pten^PEΔ/Δ^* prostates for assessment of immune cell subsets. Panel cell surface markers: cytokeratin (CK)/CD3/CD4/CD8/CD45/PD-1/DAPI. Scale bars represent 50µm. b) Quantification of tumor (left) or stroma (right) infiltrating CD3^+^ T cells reveals no statistical difference.

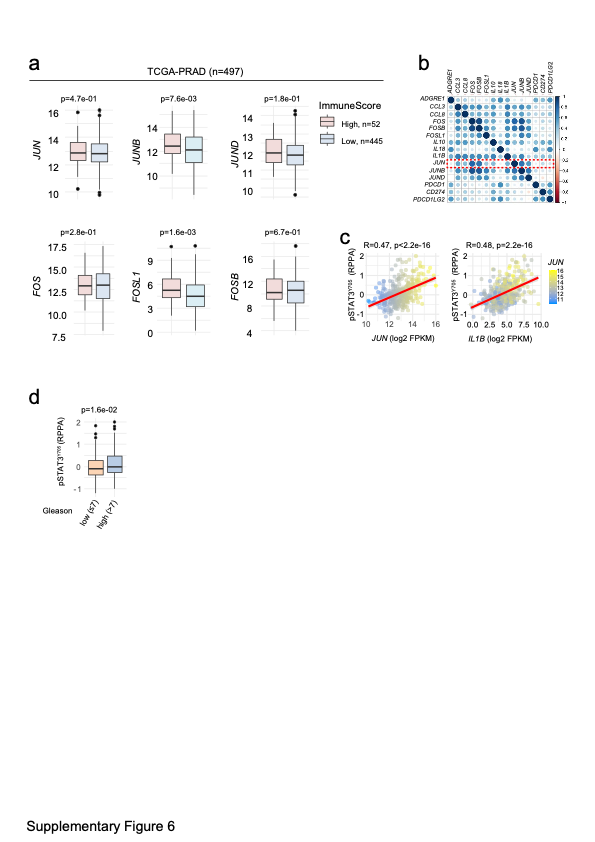

**Supplementary Figure 6: Expression of AP-1 factors *JUNB* and *FOSL1* correlate with ImmuneScore in human PCa.** a) Expression levels of indicated genes in human PCa specimens (TCGA-PRAD [40], n=497) ranked by the tumor’s ImmuneScore in high (n=52) and low (n=445). Expression of *JUNB* and *FOSL1*, but not of *JUN, JUND, FOS* and *FOSB* were significantly associated with ImmuneScore. Significance was determined by an unpaired, two-sided t-test. b) Correlation map of JUN family members and candidate mediators of immune cell attraction. The strength of correlation is color coded. c) Correlation of *JUN* (left) and *IL1B* (right) expression to amount of phosphorylated STAT3 (pSTAT3^Y705^) in the TCGA-PRAD [40] cohort (n=352). d) Box plot representing reduced levels of pSTAT3 activation (p=1.6e-02) in high risk PCa of Gleason scores >7 (range 8-10) compared to low risk (Gleason scores ≤ 7).
