## Supplementary Methods for "JUN mediates senescence and immune cell recruitment to prevent prostate cancer progression"

**RNA isolation from prostate tissue**

25 mg of RNAlater (Sigma-Aldrich) treated snap-frozen prostate tissue was homogenized in TRI reagent (Merck) with a T10 homogenizer (IKA) according to manufacturer’s instructions. The aqueous, RNA containing phase mixed with 70% EtOH was transferred to a ReliaPrep Minicolumn (Promega) and a DNAseI digest was performed directly on the column following the manufacturer’s protocol. The DNAseI digested, cleaned RNA was eluted in 25 μl nuclease-free H_2_O and the quality control was performed on the 4200 Tapestation (Agilent). The concentration was determined on a DS-11 FX+ nanophotometer (DeNovix).

**Gene set-enrichment analyses and expression analyses**

Gene set-enrichment analysis (GSEA) and single sample GSEA were performed using the stand-alone software tool (GSEA v4.3.2) or latest R script GSEA-R v1.2 and self-defined gene signatures or defined signatures as provided by the Broad Institute MSigDB, version 6.1.1 [29]. We utilized the H, C2 (hallmark gene sets, human collections), MH (mouse-ortholog hallmark gene sets) and M5 (ontology gene sets) or single gene signatures (“Inflammatory response”, “Neutrophil migration/chemotaxis”) and SenMayo (mouse) signature [30]. Genes represented in the “InnateDB_core” signature were derived from “InnateDB: Systems Biology of the Innate Immune Response” [31]. All gene signatures are available in Supplementary Table 5. For representation of “Top differentially regulated pathways” (Fig. 3d), we performed GSEA using H and C2 and selected the top enriched (FDR≤0.05) signaling pathways of comparisons *wt* versus *Pten^PEΔ/Δ^* and *Pten^PEΔ/Δ^* versus *Jun^PEΔ/Δ^;Pten^PEΔ/Δ^.* For gene ontology (GO)-term analysis of DEGs among *Pten^PEΔ/Δ^* versus *Jun^PEΔ/Δ^;Pten^PEΔ/Δ^* (Fig. 3g), we used signature M5. The ggplot2 R package was used for heat map and bubble chart-based representation.

**Statistical analysis of RNA sequencing data**

Box plots represent data of n≥3 samples (stated) and show median (center line), the upper and lower quartiles (the box), and the range of the data (the whiskers), including outliers. Significance was determined by an unpaired, two-sided t-test using R. Kaplan-Meier survival plots using the KMplot tool (<http://kmplot.com/private/>) were determined by cox regression analysis and statistical significance (p-value) was calculated by logrank test. We performed survival analysis using the SurvExpress webtool [32] as previously described [33]. Generally, the significance level of differences between groups was determined by two-tailed unpaired Student’s t-tests for two groups or ordinary one-way ANOVA. In GSEA analyses, only processes with p-values corrected for multiple testing (FDR-adjusted p-value <0.05) were considered significantly regulated. Expression data as determined by transcriptome profiling are represented as bar graphs with individual data points.

**Scanning**

Stained slides were scanned with a PANNORAMIC Scan II from 3DHISTECH, using the following parameters: Objective type: 20x; Output resolution: 49x native; Multilayer mode: extended focus, 7 levels, step size 1 µm; Compression: JPG; Bit depth: 8-bit; stitching enabled; and saved as MRXS files. Representative pictures for figures were exported using the snapshot function of CaseViewer (Build 2.4.0.119028).

**Multiplex immunohistochemistry**

Mouse prostate samples were stained with multiplex IHC and analysed by multispectral imaging. A panel of 6 fluorescent markers plus DAPI as a nuclear stain were used to detect the epitopes of CD3 (Abcam, clone SP162), CD4 (Abcam, clone EPR19514) CD8 (Cell Signaling, clone D4W2Z), CD45 (Abcam, clone EPR20033), PD-1 (Cell Signaling, clone D7D5W) and pan-Cytokeratin (Agilent Technologies, clone AE1/AE3). The staining of all slides was performed with autostainer system Bond RX (Leica Biosystems Inc.). The slides were then scanned with the Vectra® 3 (Akoya Biosystems; software version 3.0.7) microscope. Whole-slide scans were taken at 4x magnification to define regions of interest (whole tissue area) to be scanned in higher resolution using Phenochart software, version 1.0.12. Multispectral images of defined areas (whole tissue) were recorded with 20x magnification, resulting in one image color channel for each stained antibody. Images were processed with inForm software (Akoya Biosystems; software version 2.4.10), including spectral unmixing and removal of autofluorescence. Multispectral images were evaluated using HALO® Image Analysis Platform (Indica Labs). Single recorded images at 20x magnification are stitched together into a continuous field of view of the whole tissue. Individual cells were then identified using the DAPI nucleus staining by setting a threshold for nucleus size, roundness and signal intensity. For the 6 fluorescently labeled markers, positivity thresholds were set according to the staining intensity.

**Luminex cytokine array**

Snap frozen prostate tissue was homogenized in 500 µl Schindler’s lysis buffer [37] using a 1600 MiniG® tissue homogenizer and steel beads for 60 seconds at 1500 rpm. The homogenized solution was centrifuged for 30 minutes at 21000 xg and 4°C, after which the supernatant was transferred and measured for its protein concentration. 100 µg of protein were used to measure cytokine concentration using the ProcartaPlex Mouse Basic Kit (Invitrogen™, EPX010-20440-901) in combination with the respective simplex kits for each cytokine (Invitrogen™, EPX01A-26002-901, EPX01A-20603-901, EPX01A-20607-901). Immunoassays were performed according to the manufacturer’s instructions and measured with a Bio-Plex 200 (Bio-RAD) system. Significance was determined using an ordinary one-way ANOVA with Tukey’s multiple comparisons tests for three or more groups. Graphs were created and formatted in GraphPad PRISM (version 9.5.0).
